# Plant LETM1 homologs are required for fungus-induced antibiotic resistance and biostimulation

**DOI:** 10.64898/2026.04.13.718132

**Authors:** Bradley R. Dotson, Saritha Panthapulakkal Narayanan, Sara Behnamian, Sakthivel Kailasam, Mihir Shah, Thomas Kraft, John Schmidt, Tobias Ekblad, Elisabeth Veeckman, Kenneth Fredlund, Laura Grenville-Briggs, Allan G. Rasmusson

## Abstract

Our findings confirm that *CIRA15A* / *LEUCINE ZIPPER-EF-HAND CONTAINING TRANSMEMBRANE PROTEIN1* (*LETM1*) is critical for *Trichoderma*-induced growth biostimulation in sugar beet and the Cellulase-Induced Resistance to Alamethicin (CIRA) response in *Arabidopsis*. Notably, this plant homolog of a gene associated with human disease plays a vital role in both defense against *Trichoderma* antimicrobial peptides and the biostimulation of plant growth. We identified *AtCIRA15A*/*LETM1* and *AtCIRA15B*/*LETM2* as key genetic determinants of CIRA through *Arabidopsis* analysis and comparative studies of sugar beet inbred lines. *BvLETM1* allelic variations correlated with differential biostimulation responses, and complementation confirmed functional *LETM1* alleles restore CIRA in *Arabidopsis* mutants. These findings highlight LETM1 as a crucial factor in *Trichoderma*-plant interactions, with potential applications in breeding for enhanced microbial-induced plant biostimulation and agricultural productivity.

## Introduction

Plant growth stimulation and disease suppression by beneficial microbes such as *Trichoderma* vary widely among plant genotypes, yet the plant genetic factors controlling this variation remain largely unknown (Shoresh & Harman 2008; Tucci, Ruocco, De Masi, De Palma & Lorito 2011; Schmidt *et al*. 2020). These responses require plant cellular processes that translate microbial signals into physiological changes. The crop sugar beet (*Beta vulgaris subsp. vulgaris*) is the major source of sugar production in temperate zones and is a robust part of crop rotation systems in modern agriculture (Stevanato *et al*. 2019). *Trichoderma* species have been reported to promote growth and protect against several diseases in sugar beet (Abada 1994; Galletti, Burzi, Cerato, Marinello & Sala 2008; Moayedi 2010; Kappel, Kosa & Gruber 2022; Guzmán-Guzmán, Etesami & Santoyo 2025), yet plant breeding for successful biostimulation and/or biocontrol by *Trichoderma* has not yet been widely explored (Schmidt *et al*. 2020). We previously reported that within a small but diverse group of inbred sugar beet lines, there was growth stimulation in less than half of the lines tested with the commercially available *Trichoderma afroharzianum* strain T22 (Taf), and at least one breeding line demonstrated a negative growth effect after Taf inoculation despite seemingly similar root colonization (Schmidt *et al*. 2020). This variability may reflect differences in communication between the *Trichoderma* strain and the host plant.

Beneficial effects of *Trichoderma* derive from their capacity for the production of antimicrobial and defense-related compounds (Vinale *et al*. 2009; Alfiky & Weisskopf 2021; Pacheco-Trejo *et al*. 2022). However, some *Trichoderma* defense compounds can also harm plant cells, such as the specialized antibiotic peptides called peptaibols are toxic to plants such as *Arabidopsis* (Matic, Geisler, Moller, Widell & Rasmusson 2005; Rippa, Adenier, Derbaly & Béven 2007). Peptaibol peptides are amphiphilic and bind to membranes with a membrane potential when approaching from the net positively charge side, where they aggregate to form pores that disrupt membrane integrity, leading to the leakage of cellular ions and metabolites, and ultimately, cell death (Shi *et al*. 2016; Dotson *et al*. 2018; Panthapulakkal Narayanan *et al*. 2026). Interestingly, *Trichoderma* also secrete plant cell wall-degrading cellulases, partial degradation of the plant cell wall promotes modification of the plant plasma membrane, including remodeling of phospholipid composition, to become resistant to alamethicin (Aidemark *et al*. 2010). This inducible antibiotic resistance process is known as Cellulase-Induced Resistance to Alamethicin or the CIRA response (Dotson *et al*. 2018; Panthapulakkal Narayanan *et al*. 2026). Genes essential for the CIRA response have recently been identified; *AtCIRA12* and *AtCIRA13*, which encode a phosphatidylserine decarboxylase (*PSD3*) and a phospholipase (*PLDζ2*), respectively, are both required for the lipid remodeling of the plasma membrane that takes place during the CIRA response (Panthapulakkal Narayanan *et al*. 2026). The existence of an inducible symbiotic system in plants can potentially explain the large variability between plant species and cultivars in response to the same *Trichoderma* strain (Shoresh & Harman 2008; Tucci *et al*. 2011; Schmidt *et al*. 2020). This observation prompted investigation into *Arabidopsis* genes essential for CIRA and to assay whether these *Trichoderma*-associated homologous genes in sugar beet contribute to biostimulation by Taf.

LEUCINE ZIPPER EF-HAND CONTAINING TRANSMEMBRANE PROTEIN 1 (LETM1) is an integral protein in the inner membrane of mitochondria in all eukaryotes. The mammalian LETM1, and its homologue in yeast (MDM38), have been intensively studied because it is important for Ca^2+^ and/or K^+^ homeostasis in mitochondria and the cytosol (Austin & Nowikovsky 2019; Mohammed & Nowikovsky 2024). Furthermore, *HsLETM1* has been associated with the genetic disease Wolf-Hirschhorn syndrome, in which affected individuals display defects in growth, tonus, intellectual ability, cranium, and heart, and is associated with multiple cancer forms and type 2 diabetes (Li *et al*. 2019; Austin & Nowikovsky 2019; Garbincius & Elrod 2022). HsLETM1 has been studied in isolation and reported to be a Ca^2+^/H^+^ antiporter (Jiang, Zhao & Clapham 2009; Tsai, Jiang, Zhao, Clapham & Miller 2014; Shao *et al*. 2016), which may also be involved in Ca^2+^ export (Garbincius & Elrod 2022), but was more recently shown to be an interaction partner to the Ca^2+^/H^+^ antiporter TMBIM5, yet a more exact function of LETM1 is not known (Austin *et al*. 2022). Mammalian LETM1 may additionally be involved in K^+^/H^+^ exchange, as reported for yeast MDM38, and the molecular function of LETM1 remains debated (Austin & Nowikovsky 2019; Austin *et al*. 2022). Studies of the *Arabidopsis* homologs to *HsLETM1*, *AtCIRA15A/LETM1* and *AtCIRA15B/LETM2*, have revealed that homozygous mutants of either showed no growth phenotype and double homozygous mutants were embryo-lethal (Zhang *et al*. 2012). A viable homozygous/hemizygous double mutant displayed growth reduction during early development, reduced mitochondrial translation and starch accumulation by an unknown mechanism (Zhang *et al*. 2012). It has been suggested that the LETM1 proteins in plants, like in some other eukaryotes, modify homeostasis of Ca^2+^ in both the cytosol and the mitochondrion (Carraretto *et al*. 2016), but this has not been experimentally investigated.

Here we identify the *Arabidopsis CIRA15A / LEUCINE ZIPPER EF-HAND CONTAINING TRANSMEMBRANE PROTEIN* genes *AtCIRA15A/LETM1 and AtCIRA15B/LETM2* and their ortholog in sugar beet (*BvCIRA15A/LETM1*) as essential components for plant defense against a fungal antibiotic. They are also significantly associated with Taf-induced plant growth stimulation. These genes are thus potential markers for trait selection for more efficient biostimulation by *Trichoderma* in future crop cultivars.

## Methods and Materials

### Preparation of seedling cultures

Wild-type (Col-0) and T-DNA mutant seeds of *Arabidopsis thaliana* (Lehle Seeds, Round Rock, TX, USA) were surface sterilized with 70% (v/v) ethanol for 1 min, followed by 2.5% (w/v) sodium hypochlorite containing 0.1% Tween 20 for 1 min, and washed five times with distilled water. Seeds were then transferred either to 3.5 cm Petri dishes for cultivation in H₂O or to 96-well polypropylene plates with lids (Greiner Bio-One, BioNordika, Stockholm, Sweden) containing 100 μl sterile H₂O and ∼5-10 seeds per well. After 2-3 d of stratification at 4°C, seedlings were transferred to a growth room (16 h light, 24°C). Seedlings grown on Petri dishes were used for fluorescence microscopy assays after 4-10 d, whereas seedlings grown in 96-well plates were used for CIRA assays after 5 d.

Dried sugar beet fruits containing seeds were transferred to separate tea strainers and soaked in 70% ethanol for 1 min, then transferred to 2.5% (w/v) sodium hypochlorite for 1 min and washed three times with H2O. The fruits were then submerged in 60°C tap water for 10 min and placed in a beaker under a constant flow of tap water for ∼16 h. Seeds from each line were transferred onto pleated filter paper inside polyethylene containers, moistened, and allowed to germinate under 16 h light at 22-24°C.

### CIRA assays

*Arabidopsis* seedlings grown in 96-well polypropylene plates were subjected to mock or 1% *Trichoderma* cellulase (“Onozuka RS”; Duchefa, Haarlem, Netherlands) treatment at pH 5.0 with gentle shaking for 2 h at 20°C. The solution was removed, and the seedlings were washed three times with H₂O, after which the wells were replenished with 100 µL of either H₂O or 20 µg/mL alamethicin (Sigma-Aldrich, St. Louis, MO, USA) and treated for 10 min with gentle rocking at room temperature. During the final minute of the 10-min treatment the solution was supplemented with 1 µL of 0.15 mM propidium iodide (PI), a membrane-impermeable DNA stain. The solution was then removed and saved for conductivity measurements, and the seedlings were replenished with 100 µL diH2O and subjected to fluorescence microscopy.

Sugar beet seeds used for CIRA assays were sterilized, germinated, and grown in sterile sand. After 7 d, seedlings were removed from the sand and placed in 15 mL Falcon tubes (Sigma-Aldrich, St. Louis, MO, USA). Three-to five-day-old seedlings from each line were subjected to control treatment or cellulase treatment (1% *T. viride* cellulase for 2 h) followed by 10 min treatment with 20 µg/mL alamethicin. Seedlings were then stained with 1 µM PI for 1 min prior to analysis.

### Conductivity measurements

Conductivity measurements of the CIRA assay were performed as previously described in Panthapulakkal Narayanan *et. al.* (2026). Solutions analyzed for ion leakage (µS/cm) were measured using a CDM230 conductivity meter (Radiometer Analytical, Brønshøj, Denmark) at 20°C (Panthapulakkal Narayanan *et al*. 2026). Resulting measurements from at least three biological replicates were normalized to the number of germinated seedlings.

For lipid perturbation experiments, lysophosphatidylserine (LPS; Avanti Polar Lipids, product no. 858143) was added during the cellulase treatment (Panthapulakkal Narayanan *et al*. 2026). Attempts to quantify the CIRA response through electrolyte leakage in sugar beet were confounded by background metabolite leakage, leading to inconclusive results (data not shown).

### Fluorescence microscopy

Fluorescence microscopy of *A. thaliana* and sugar beet seedlings was performed using a G-2A filter set (excitation 510–560 nm, emission >590 nm) coupled to a Nikon Optiphot-2 microscope (Nikon Corporation, Tokyo, Japan). Images were collected using an Olympus DP-70 digital camera (Olympus Optical, Tokyo, Japan). Exposure times were determined using control roots treated with 20 µg/mL alamethicin and applied consistently to all images within each experiment. Post-processing of images was performed using ImageJ software (NIH) (Magelhaes, Ram & Abramoff 2004).

### Trichoderma culturing

*Trichoderma afroharzianum* strain T22 (ATCC 20847) was cultured on potato dextrose agar (PDA) (Sigma-Aldrich, St. Louis MO, US) at 20℃ for 3 weeks with 16/8-hour illumination. Mature spores were harvested with sterile deionized H_2_O filtration through sterile cotton ball mesh. The spore stock culture was stored at 4℃ until use. The culture’s colony-forming units (CFU) were determined by serial dilution plating on PDA plates.

### Sugar beet soil growth

Seedlings with fully emerged cotyledons were transferred into a 3 × 5 × 18 cm column tube system containing potting soil (Gröna linjen, SW Horto AB). Plants were grown in a greenhouse with supplemental metal halide lighting (Osram Powerstar HQI-BT; 16 h photoperiod; approximately 150 µmol m⁻² s⁻¹ at ∼20°C). Two days after transplantation each seedling was treated with either 500 µL H₂O or 500 µL H₂O containing 10,000 CFU of Taf. Seedlings were then grown for an additional 7 d before harvest.

During harvesting, potting soil was carefully removed, the primary root length was measured, and shoot and root tissues were separated into pre-weighed aluminum foil packets. Samples were dried at 80°C for approximately 2 d. Shoot and root dry weights were then combined to calculate total dry weight. Additionally, root dry weight was divided by primary root length to estimate lateral root dry weight per primary root length.

These measurements (Total, Shoot, Root, Lateral Root Weight, and Primary Root Length) were log₂-transformed. For each set of Taf-treated seedlings, log₂-transformed values were normalized to the untreated control within each experiment by subtracting the mean log₂ value of the control. The shoot-to-root ratio was calculated by subtracting the log₂ root dry weight from the log₂ shoot dry weight.

Outliers were removed according to Tukey’s fences for identifying outliers (Tukey 1949). Data were then tested for significance using Student’s t-test with false discovery rate (FDR) correction (*q-*value of 0.05) (Benjamini, Drai, Elmer, Kafkafi & Golani 2001).

### Barley soil growth

Barley (*Hordeum vulgare* L.) lines Barke, Bonus, Morex, and FT880 (Leibniz Institute of Plant Genetics and Crop Plant Research, IPK Genebank, Germany) were directly sown and treated in a similar manner to sugar beet seedlings.

### DNA extraction and Polymerase Chain Reaction (PCR)

Plant tissue was ground in a 1.5 mL microcentrifuge tube using liquid nitrogen and a plastic pestle. DNA extraction was then performed according to the manufacturer’s protocol using the DNeasy Plant Mini Kit (Qiagen, Hilden, Germany; ref. 69106). Resulting DNA extracts were stored at −20°C until further use.

PCR was performed using the DreamTaq Green PCR kit (Thermo Fisher Scientific, Waltham, MA, USA) according to the manufacturer’s instructions. PCR products were separated on 1.5% agarose gels prepared in 0.5× TAE buffer and electrophoresed at 95 V before imaging.

### Sugar beet genome resequencing and variant calling

Whole-genome resequencing was performed on 25 sugar beet (*Beta vulgaris*) inbred accessions. Whole-genome DNA data consisted of seven accessions sequenced using the Illumina HiSeq 2500 platform (BGI, Beijing, China) and 18 accessions sequenced using the Illumina HiSeq 4500 platform (BGI, Poland). All accessions were sequenced as 150 bp paired-end reads. Quality control was performed using NGS QC Toolkit (Patel & Jain 2012). Adapter removal and read trimming were performed using Trimmomatic (Bolger, Lohse & Usadel 2014) and read quality was assessed using FastQC (www.bioinformatics.babraham.ac.uk/projects/fastqc).

An initial set of inbred lines (A, B, D, F, G, H, and I) was mapped to the *Beta vulgaris* reference genome RefBeet 1.5 (Dohm *et al*. 2014; Minoche *et al*. 2015), to determine the candidate marker region. Subsequently, sequence data from all inbred lines were mapped to the EL10.2 reference genome (McGrath *et al*. 2023) using the BWA aligner (Li & Durbin 2009). Alignment files were further processed using Picard Tools (https://broadinstitute.github.io/picard/).

Genotypes were called using HaplotypeCaller from the Genome Analysis Toolkit (GATK) (McKenna *et al*. 2010) with a minimum base quality score of 20. HaplotypeCaller produced a variant call file (VCF) for each inbred line, which were subsequently combined using GATK CombineGVCFs. Variant polymorphisms within the candidate marker region were analyzed to determine similarity of gene loci to the initial inbred line set using Geneious (Kearse *et al*. 2012)

### Sanger sequencing and mutation conformation

Allele verification for selected loci in *Arabidopsis thaliana*, sugar beet, and barley was performed by PCR amplification of genomic regions containing diagnostic polymorphisms. Gene-specific primers were designed, and PCR products were analyzed on agarose gels to confirm the presence of a single amplicon (Fig. S1). PCR products were then purified and submitted for Sanger sequencing (Eurofins Scientific, Luxembourg City, Luxembourg).

### Evolutionary analysis

*CIRA15/LETM* homologs were identified using BLAST searches against NCBI databases. For identification of CIRA homologs, the genomes of *Arabidopsis thaliana* (Col-0), *Beta vulgaris* subsp. *vulgaris*, *Chenopodium quinoa*, and *Oryza sativa* (japonica group) were used to establish clade boundaries. After identification of suitable homolog candidates, annotated mRNA coding sequences were translated and aligned using the BLOSUM62 substitution matrix in MUSCLE (Edgar 2004), All phylogenetic analyses were based on coding sequence comparisons.

Subsequent phylogenetic analysis was performed in Geneious (Kearse *et al*. 2012) using the maximum likelihood algorithm implemented in PhyML (Guindon *et al*. 2010) with the HKY85 substitution model and 1000 bootstrap replicates. In the larger LETM1-like phylogeny, regions of low sequence conservation (amino acids 550–581 of BvCIRA15A/LETM1) were masked prior to tree construction (Fig. S2). Homolog candidates were further evaluated based on sequence similarity across kingdoms. The complete list of sequences used in the analysis is provided in Fig. S3. To maintain balanced representation within the phylogeny, selected species were removed to reduce overrepresentation of taxa within Plantae.

### *Agrobacterium*-mediated transformation of *Arabidopsis*

Coding sequences (CDS) of alleles of *BvCIRA15A/LETM1.I/G* and *BvCIRA15A/LETM1.F* and *AtCIRA15A/LETM1* (At3g59820) were inserted to in a construct containing the *AtCIRA15A/LETM1* promoter (-1000bp) with 5’ and 3’ untranslated regions (UTRs), upstream of a NOS terminator. Constructs were designed *in silico* and synthesized (Genescript, Piscataway NJ, USA) (Fig. S4-S6).

The resulting constructs were cloned into the Agrobacterium binary vector pC1300, containing hygromycin resistance as a plant transformation marker. Constructs were verified by PCR and restriction enzyme digestion before transformation.

The three vectors were introduced into *Arabidopsis thaliana* Col-0, *cira15a-1*, *cira15b-1*, and *cira15b-2* backgrounds via *Agrobacterium*-mediated floral dip transformation (Clough & Bent 1998). Transformants were selected on hygromycin-containing medium. For each construct–background combination, three independent transgenic lines were selected for subsequent CIRA assays.

## Results

*Arabidopsis CIRA15/LETM* gene paralogs are each essential for cellulase-induced alamethicin resistance (CIRA)

Antibiotic peptide alamethicin permeabilization of a plasma membrane is detected by the intracellular accumulation of the membrane-impermeable nucleic acid-intercalating probe propidium iodide (PI) (Fig. 1A). In wild type, plasma membrane permeabilization by alamethicin can be counteracted by pretreatment of the roots with *Trichoderma* cellulases with the CIRA response (Fig. 1A) (Dotson *et al*. 2018; Panthapulakkal Narayanan *et al*. 2026). To identify genes that are essential for CIRA in *Arabidopsis*, we performed a genetic screen of a collection of T-DNA mutants of *Arabidopsis thaliana* Col-0. We identified that mutations in *AtCIRA15A/LETM1* (*At3G59820*) or *AtCIRA15B/LETM2* (*At1G65540*), *CELLULASE INDUCED RESISTANCE TO ALAMETHICIN 15A* and *15B,* were the underlying causes of *cira15a* and *cira15b* mutants, respectively (Fig. 1A, Fig. S7A, and S7B). Both *Atcira15a/letm1* and *Atcira15b/letm2* mutants were deficient in the CIRA response observed in the root tip extension zone. For each mutant, additional mutant alleles confirmed the CIRA phenotype. Without cellulase treatment, wild type (Col-0) and all mutant lines displayed similar alamethicin-dependent propidium iodide (PI) staining. In the absence of alamethicin, neither control nor cellulase-treated seedlings displayed PI staining. After cellulase treatment, Col-0 roots showed no PI staining despite alamethicin presence, but all *cira15a* and *cira15b* mutant lines displayed staining at the root tip, similar to the treatment with alamethicin alone (Fig. 1A, Fig. S7B). The loss of CIRA function in *cira15a* and *cira15b* mutant lines was confirmed through quantification of leaked electrolytes into the supernatant due to alamethicin (Fig. 1B). We found that alamethicin treatment alone led to a similar increased medium conductivity in Col-0 and the *cira15a* and *cira15b* mutant lines, indicating that all lines were equally susceptible to alamethicin-dependent loss of electrolytes. After pretreatment with cellulase, the loss of electrolytes was strongly decreased in Col-0 seedlings, the CIRA response. By comparison, the *cira15a* and *cira15b* seedlings released similar amounts of electrolytes irrespective of cellulase pretreatment, thus lacked a CIRA response (Fig. 1B). These two approaches demonstrated that the genes *AtCIRA15A/LETM1* and *AtCIRA15B/LETM2* each participate in and are essential for the CIRA process.

**Figure 1.**
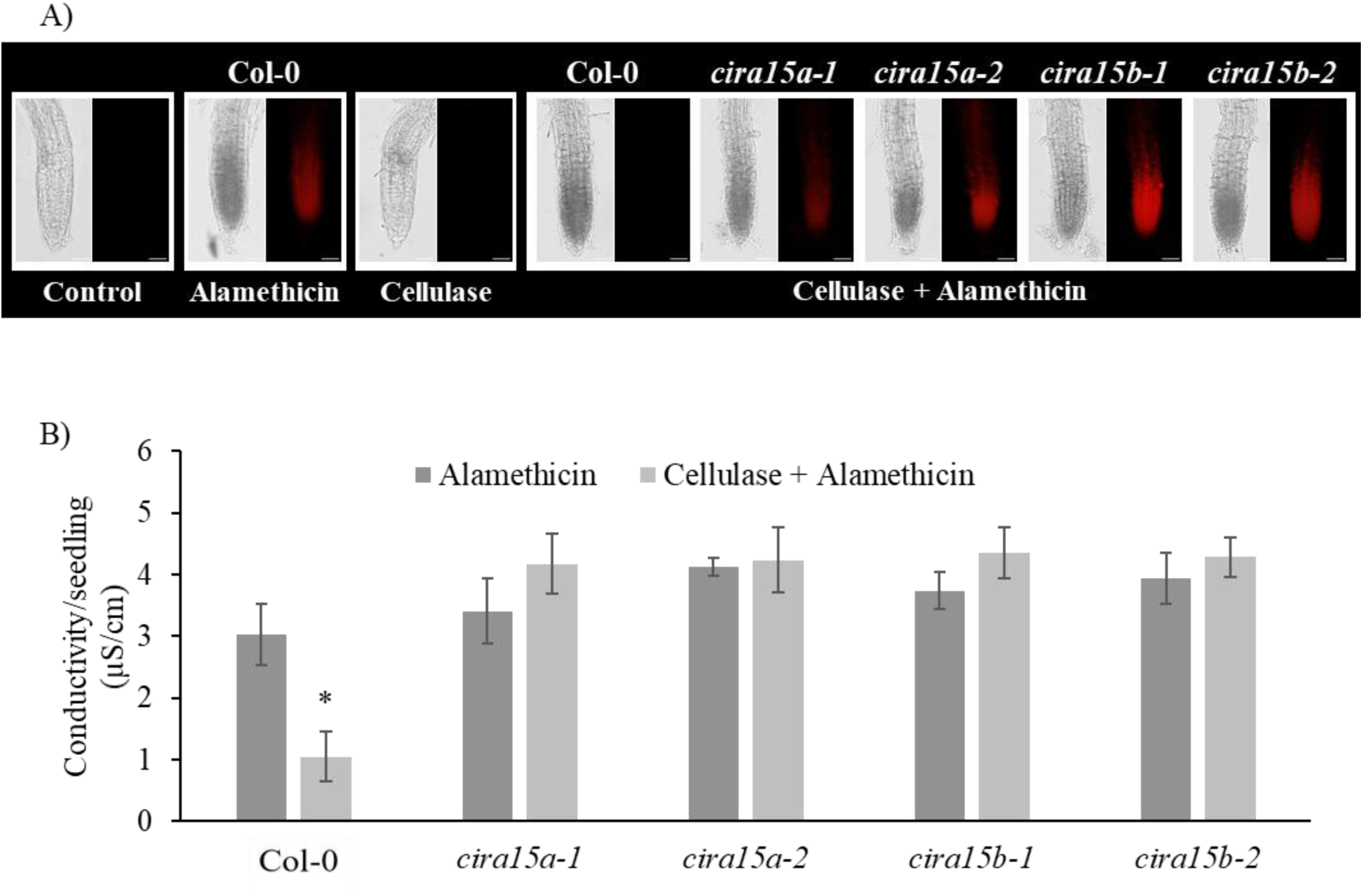
*AtCIRA15A/LETM1* and *AtCIRA15B/LETM2* are essential for cellulase-induced resistance to the peptaibol alamethicin. A) Representative *Arabidopsis* seedling roots from wild type Col-0, and mutant lines were treated with *Trichoderma* cellulase to induce resistance to alamethicin and analysed by bright field and fluorescent microscopy. Red fluorescence indicates penetration of alamethicin into the plasma membrane, causing influx of propidium iodine. Both wild type and mutants displayed similar results for control, alamethicin and cellulase treatment. Bars represent 50 µm. B) Alamethicin-induced ion leakage from *Arabidopsis* seedling roots wild type Col-0 and *Atcira15* mutants was measured by conductimetry. Alamethicin treatment leads to electrolyte loss from the cells, but after pre-treatment with *Trichoderma* cellulases leakage is reduced due to the CIRA response. Bars represent standard error, and asterisks denote significant differences between cellulase pretreated and control using an FDR *q*-value of 0.05 (*n* = 3 biological replicates).

*Arabidopsis AtCIRA15/LETM* gene paralogs belong to the highly conserved LETM family in eukaryotes *AtCIRA15A/LETM1* and *AtCIRA15B/LETM2* were identified within the family of LETM1 domain-containing proteins pfam07766 (Mistry *et al*. 2021). Analysis of LETM1 domain-containing proteins revealed conserved LETM RBD motif with a transmembrane region (Fig. S8 and S9). Additionally, several C-terminal leucine zipper repeats creating coiled-coil domains and an EF-hand Ca^2+^-binding domain were commonly observed (Fig. S9) (Natarajan, Mishra, Camara & Kwok 2021). A phylogenetic analysis of eukaryotic homologues revealed the evolution of the EF-hand domain-containing LETM1-like proteins in plants (Fig. 2). All species studied contained at least one LETM1 domain-containing protein. Apart from Ascomycota fungi and a Basidiomycota species, at least one of the LETM1 domain-containing homologs in each species also contained an identifiable EF-hand domain and multiple C-terminal leucine zippers coiled coil domains (Fig. 2, Fig. S3, and S8-S9). Some species also contained truncated forms of LETM1 domain-containing proteins, here termed LETMD-like, which lacked the C-terminal EF-hand and leucine zipper coiled coil domains (Fig. S8-S9). LETMD-like homologs showed similarity to the human *HsLETM2* and *HsLETMD* proteins, which were less conserved and notably did not rescue *HsLETM1* function (Tamai *et al*. 2008) (Fig. S3 and S9), and were excluded from the phylogeny.

**Figure 2.**
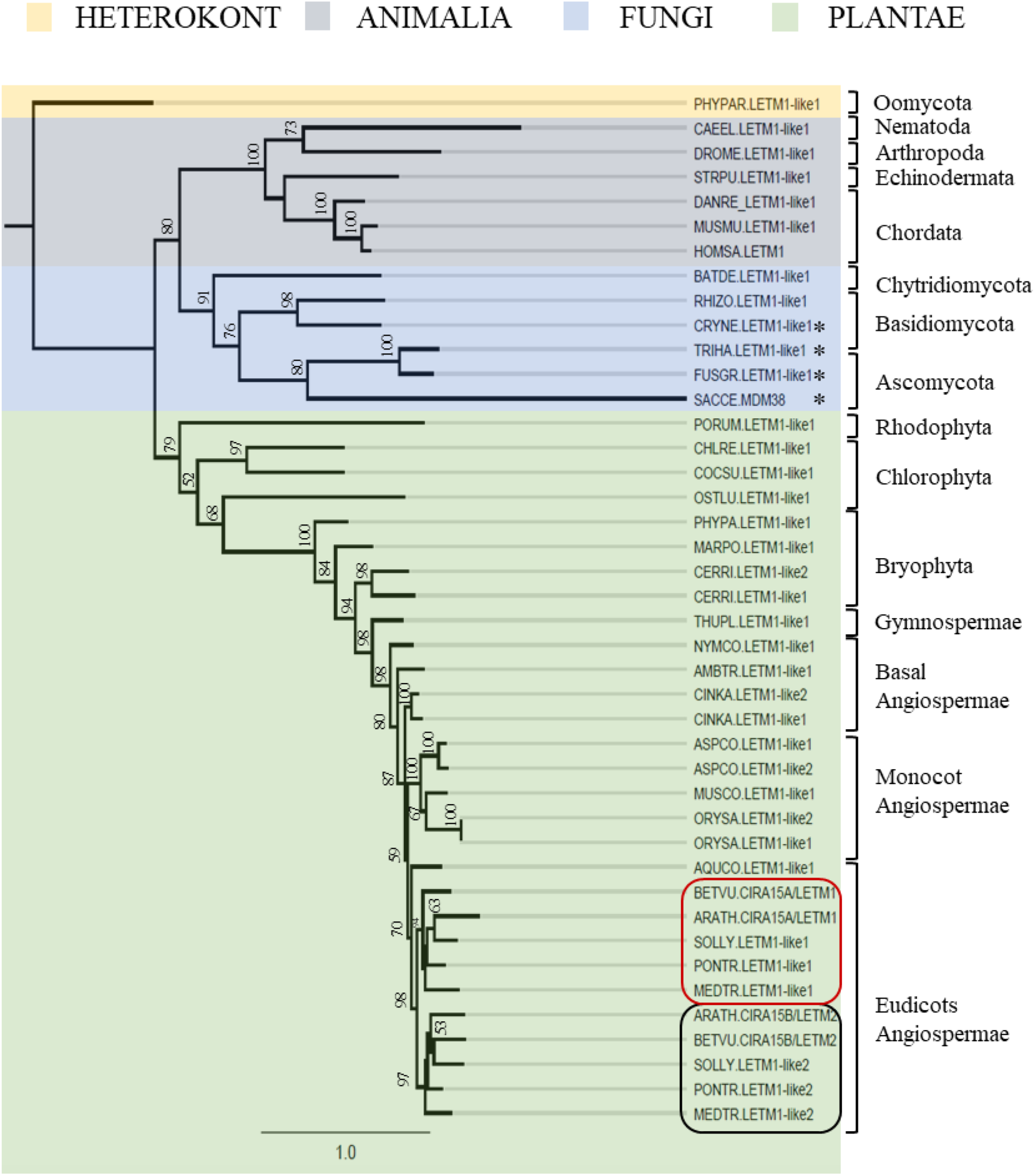
Phylogeny of LETM1-domain genes. Coding sequences for EF-hand containing LETM1 genes from Plants, Fungi and Animals were aligned and analysed by maximum likelihood using a Heterokont homolog as outgroup. LETM1-like genes that also contained EF-hand pair superfamily domains (IPR011992) or the largest LETM1-like genes were selected in each species; an asterisk denotes the absence of EF-Hand in LETM homologues within the phylogeny. Genes highlighted in red and black belong to the *CIRA15A/LETM1* and *CIRA15B/LETM2* clade respectively within the core Eudicot Angiospermae. Gene information can be found in Fig. S3. Bootstrap values are denoted where above 50. Scale represents substitution rate.

The dual paralogs of LETM1 containing EF hand domains are general in core eudicots, indicating a gene duplication in the early eudicot evolution conserved as two homologous clades, hereon renamed the CIRA15A/LETM1 clade and the CIRA15B/LETM2 clade (Fig. 2). Within the CIRA15A and CIRA15B clades, *AtCIRA15A/LETM1* correspond to the homolog in sugar beet called *BvCIRA15A/LETM1*. Similarly, the *AtCIRA15B/LETM2* gene corresponded to the sugar beet gene called *BvCIRA15B/LETM2* (Fig. 2).

A *BvCIRA15A/LETM1* polymorphism is associated with sugar beet biostimulation competency Sugar beet has displayed a genetic variation between inbred lines for the ability to be biostimulated by *Trichoderma afroharzianum* T22 (Taf) (Schmidt *et al*. 2020). A diverse group of eight sugar beet inbred lines used for breeding was previously analyzed for biostimulation by Taf in soil (Schmidt *et al*. 2020). A higher throughput method with a wider set of measured parameters was designed and compared to previously published data (Schmidt *et al*. 2020). A consistent pattern of biostimulation seven days after treatment with Taf was found (Fig. 3). Generally, the inbred lines showed a decreased shoot-to-root mass ratio in the presence of Taf. The inbred lines G and I repeatedly displayed a significant seedling stage increased growth by Taf within the phenotypes of total weight, shoot weight, root weight, and lateral root weight per length of primary root. In stark contrast, the inbred line F was significantly impaired by Taf inoculation regarding total weight, shoot weight, and primary root length.

**Figure 3.**
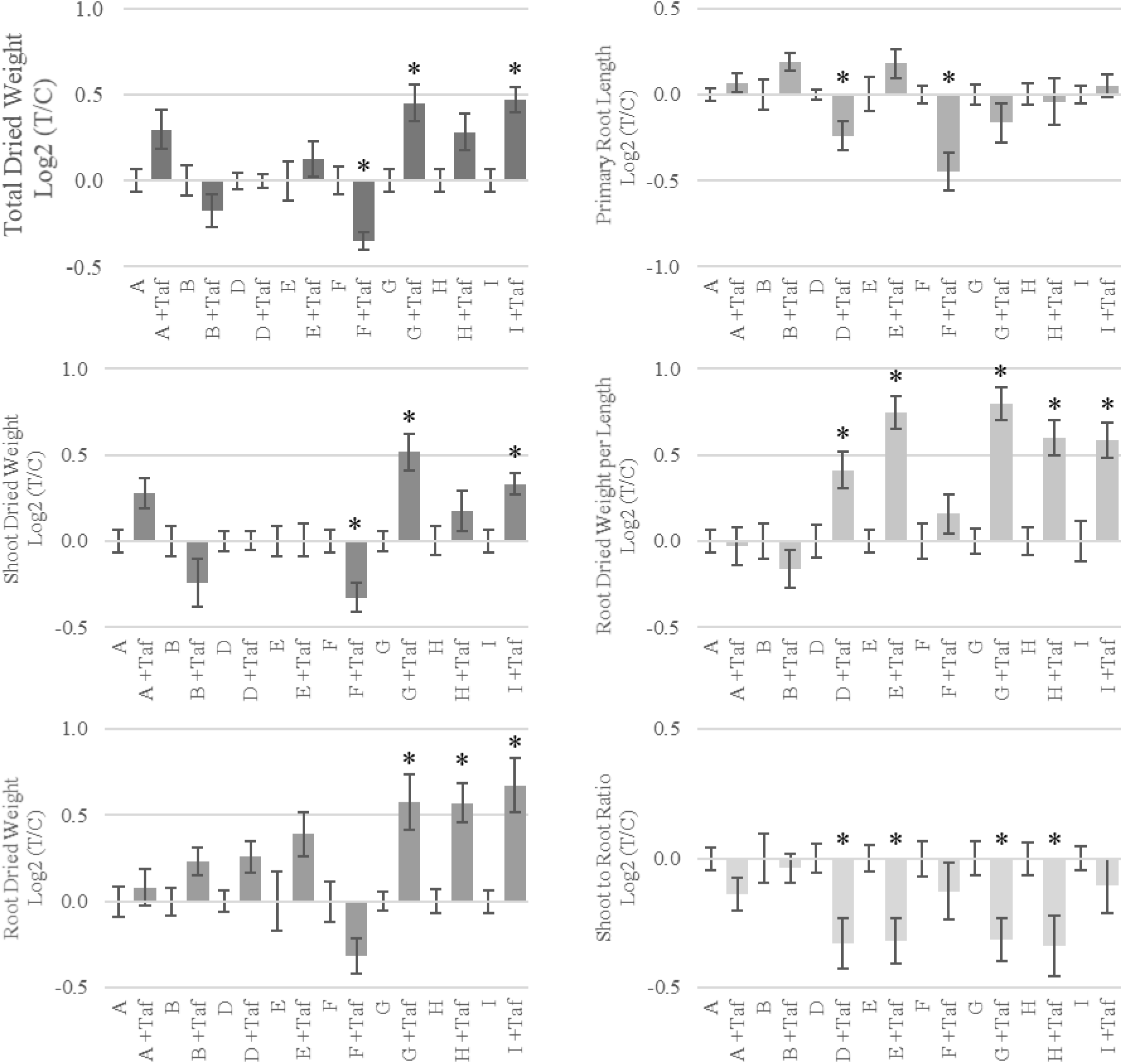
Phenotypic analysis of diverse set of inbred breeding lines. Growth statistics for eight inbred lines of *Beta vulgaris subs. vulgaris* (sugar beet). Growth comparison between lines is based on log2 transformed ratios of Taf-treated plants to untreated control plants. Bars represent standard error, and asterisks denote significant differences between treated and control using an FDR *q*-value of 0.05 (*n* = 4-8 biological replicates).

To identify the underlying genetic causes behind the differential biostimulation responses to Taf inoculation (Schmidt *et al*. 2020), the genomes of sugar beet inbred lines were sequenced and aligned to the reference genomes Refbeet 1.5 (Dohm *et al*. 2014; Minoche *et al*. 2015) for marker associations and to EL10.2 (McGrath *et al*. 2023) for sequence analysis. A candidate list of sugar beet homologues to *Arabidopsis* CIRA candidate genes and other previously *Trichoderma*-associated genes was investigated with respect to association to the phenotypic changes (Dotson, Verschut, Flärdh, Becher & Rasmusson 2026; Panthapulakkal Narayanan *et al*. 2026). The greatest difference between the inbred lines G and I to inbred line F was total weight, therefore total weight phenotypes were evaluated in an additional set of 29 sugar beet inbred lines. Markers flanking each of the candidate gene regions were identified from the available industrial breeding markers and used to classify lines as I/G or F-type for each region. A correlation analysis of the log2-transformed total dry weight ratios (Taf/control) against identified allelic markers was carried out to determine allelic differential phenotypic associations within potential CIRA homolog candidates (Fig. 4). Significant differences in total weight between additional inbred lines with marker alleles for inbred line F to marker alleles for inbred lines G and I provided an indication towards a 260 kilobase region in RefBeet1.5 on chromosome 3 between the markers IIS3758 and IIS8436, containing *BvCIRA15A/LETM1* and 11 other genes (Fig. 4A). Additional inbred lines (29) were then selected for containing variation in the regions around the IIS3758 and IIS8436 genomic markers (Fig. S10). These lines were phenotypically characterized for seedling-stage biostimulation by Taf (Fig. 4B). Inbred lines (26) with markers similar to inbred lines G and I (enclosing what is here called the I/G allele) were phenotypically significantly different from inbred lines (7) that contained markers similar to the inbred line F allele-type (Fig. 4B and Fig. S2 and S10). The I/G allele-type inbred lines displayed a specific average increase of 19% biomass upon Taf treatment (Fig. 4B). Genomic resequencing of eighteen lines with sufficient replicates identified allelic variation of the marker region possessing an I/G, F or another allele associated within the mapped region identified 4 genes of interest that contained polymorphisms unique to inbred line F and were expressed in seedlings and roots (Fig. 4C, and Fig. S11). Of the twelve loci located within the region flanked by the markers IIS3758 and IIS8436, four gene candidates matched the criteria that the loci possessed F allele-unique mutations and that the genes were expressed in seedlings and roots tissue (Fig. S11). These four candidate genes were *BVRB_3g058990*, *BVRB_3g59000*, *BVRB_3g059060* and *BvCIRA15A/LETM1* (*BVRB_3g059020*) (Fig. 5). Among the four identified genes, *BvCIRA15A/LETM1* (*BVRB_3g059020*) displayed the largest protein sequence variation between the I/G-type and F-type alleles, including two short deletions and one residue change that are unique to the F-type allele (Fig. 5, Fig. S2). A variable region containing two intron and two exons and 3’UTR region of *BvCIRA15A/LETM1* was used to positively identify a set of inbred lines for the I/G-allele and the F-allele (Fig. S12). *BVRB_3g058990, BVRB_3g05900*, and *BVRB_3g059060* had unique F allele mutations that were non-conserved or polymorphisms that were also present in other homologs from species present in phylogenetic analysis (Fig. S13-S15), indicating that these alleles were likely to function normally. Thereby, the critical locus for the early biostimulation by Taf was identified to be *BvCIRA15A/LETM1* (*BVRB_3g059020*). The I and G sugar beet lines contained the allele of *BvCIRA15A/LETM1.I/G* which possessed two similar amino acid changes (S49R and E142D) in reference to EL10.2 (McGrath *et al*. 2023), whereas sugar beet line F contained the *BvCIRA15A/LETM1A.F* allele that possessed sixteen amino acid polymorphisms of which three were unique to the F-allele (KLHPVC110(del), Q218K, and T752(fs)D*), two of which constituted substantial deletions in less conserved domains (Fig. S2).

**Figure 4.**
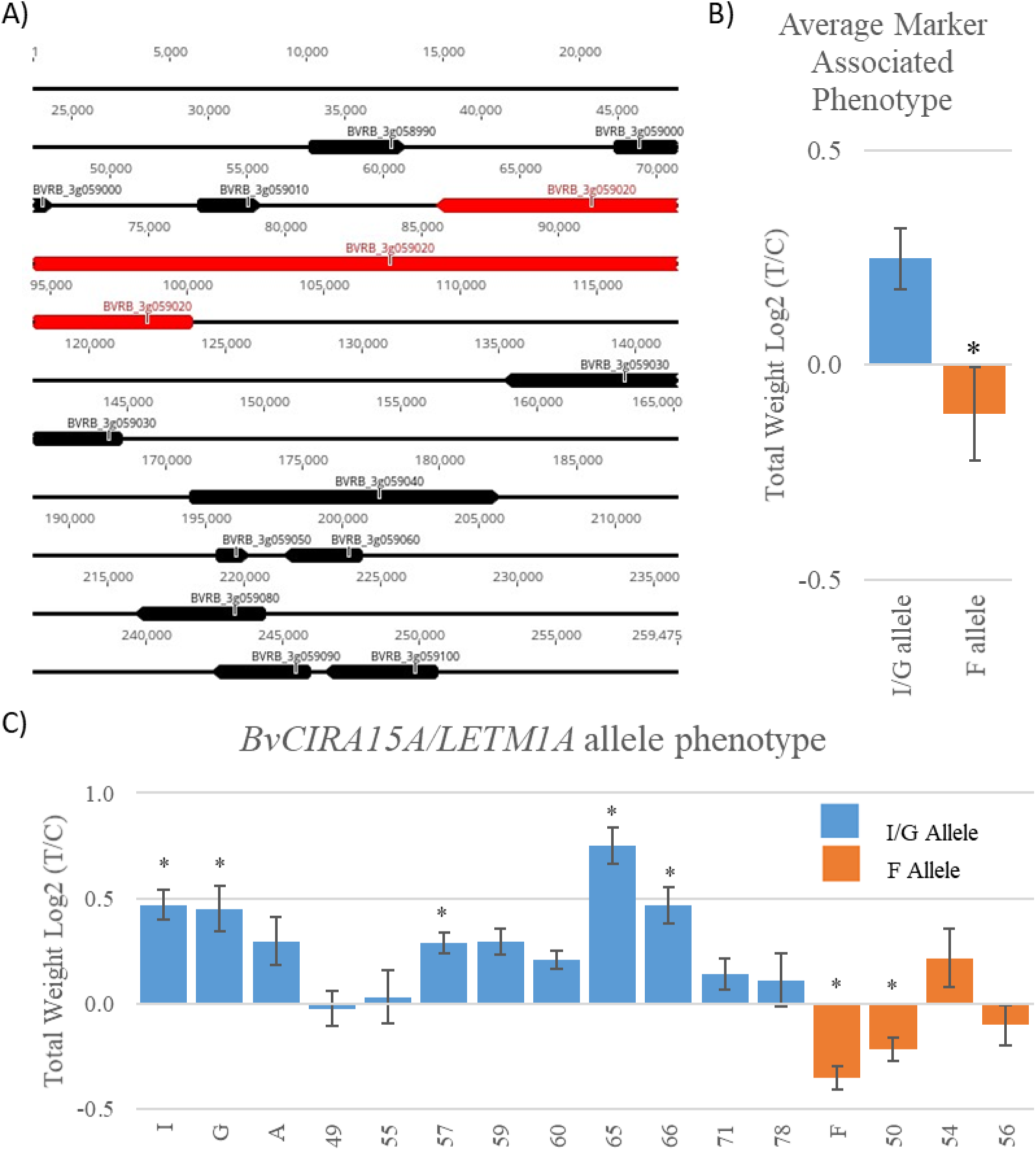
Marker identified region and allelic phenotypic analysis. A) Investigation of the defined region on chromosome 3 within Refbeet 1.5(Dohm *et al*. 2014; Minoche *et al*. 2015) that was differential between the inbred lines assayed with the markers IIS3758 and IIS8436. The gene *BvCIRA15A*/*LETM1* (colored red) was found to be less than 1 % of the chromosome length distance to the markers. B) Significant difference between the average marker associated phenotypes of I/G-type (26) and F-type (7) marker associated lines with a *p*-value of 0.017 denoted by asterisk (see Fig. S10). C) Phenotypic variation analysis for inbred lines confirmed for the I/G-type or F-type allele of the gene *BvCIRA15A/LETM1*. Bars represent standard error. Asterisk represents significant difference between control and Taf treated total weight using an FDR *q* value of 0.05 (*n* = 4biological replicates of 4 plants).

**Figure 5.**
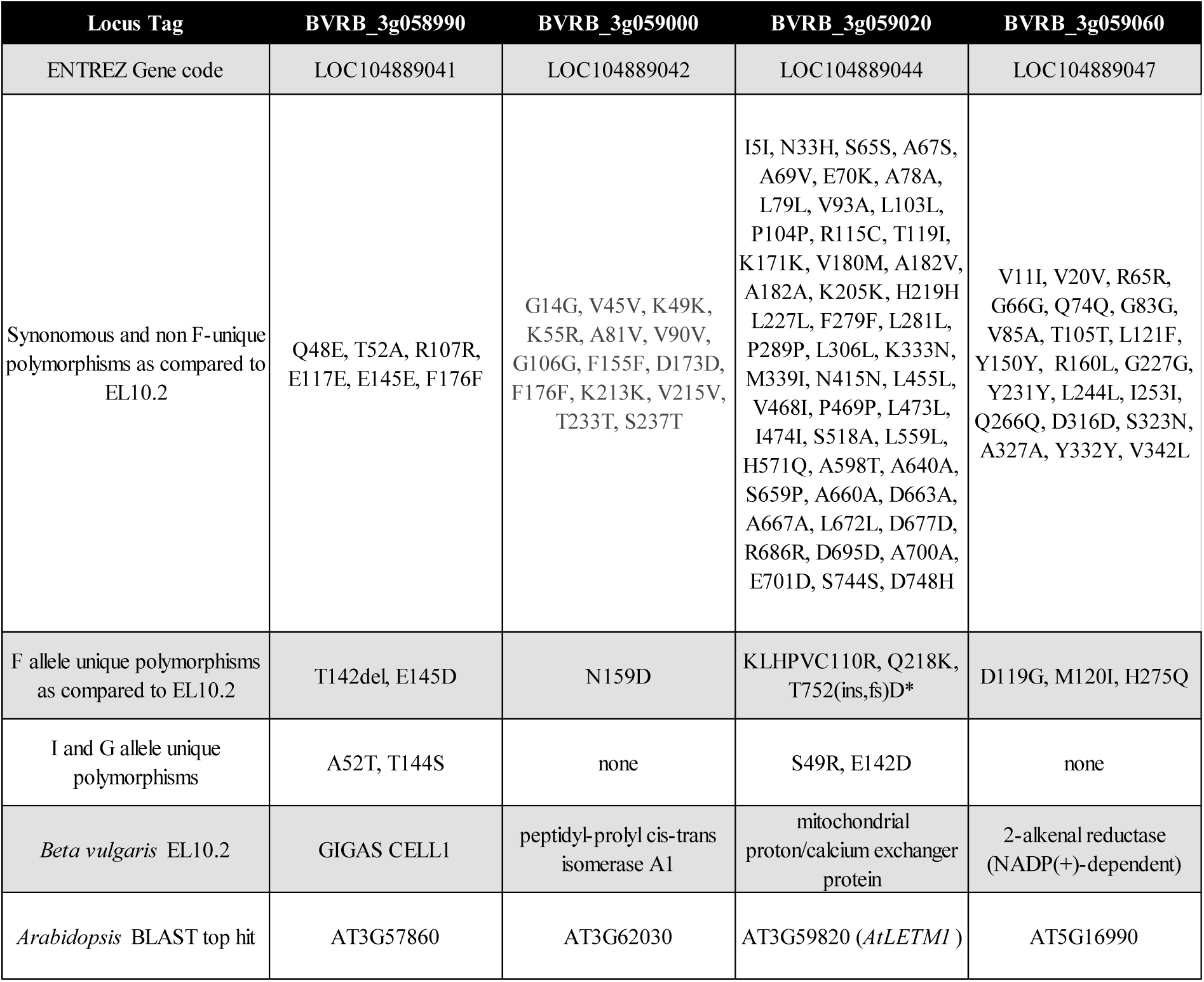
Summary of gene product information available. Summary of gene product information for loci with polymorphisms unique to F-allele identifying synonymous and non-synonymous polymorphisms, expression in sugar beet roots(Dohm *et al*. 2014; Minoche *et al*. 2015), gene description from *Beta vulgaris* EL10.2 genome annotation and top BLAST hit for *Arabidopsis thaliana*. Abbreviations of ins: insertional mutation, del: deletion mutation, fs: frameshift, and *: premature stop codon.

To investigate the potential consequences of these polymorphisms, we performed structural predictions using AlphaFold2 (Mirdita *et al*. 2022) (Fig. S16). Modelling the N-terminal part of BvCIRA15A/LETM1, including the transmembrane helix, merely produced random coils (data not shown). However, models of the C-terminal half mainly produced alpha helical structures. In homologous overlaps, these were consistent with the determined structure of the yeast MDM38 ribosome-binding domain (Lupo *et al*. 2011), except that the K^+^-binding moiety is absent in the sugar beet protein (Fig. S16). In the ultimate C-terminal part of BvCIRA15A/LETM1, the changes between *BvCIRA15A/LETM1.I/G* and the *BvCIRA15A/LETM1.F* alleles were found to shift alpha-helix positions, implying substantially different overall structures of the two allelic proteins. Therefore, the modelling suggests that the two alleles may display functional differences.

In order to identify if the sugar beet inbred lines of I and F responded differently to the CIRA assay, seedlings of the inbred lines I and F were assessed for alamethicin and CIRA responses (Fig. S17). Both lines were able to react to alamethicin and cellulase from *Trichoderma spp.* as seen from changes in PI fluorescence, though the patterns of cell permeabilization by alamethicin application differed from *Arabidopsis* (Fig. 1A, Fig. S7B). The results show that there are differences in alamethicin permeabilization between root cells of the two sugar beet lines. Line I specifically displayed a strong permeabilization in the apical meristem (Fig. S17), which could be completely counteracted by pretreatment with cellulase. In the expansion zone, a decrease in permeabilization was observed upon cellulase treatment in both lines (Fig. S17). The disparity of alamethicin susceptibility between the lines is unsurprising considering the diversity in genetic backgrounds of the inbred lines.

The absence of Taf-induced biostimulation in *A. thaliana* prompted us to examine whether this response occurs in barley (*Hordeum vulgare* L.).. The published barley pan-genome was screened for mutations predicted to result in premature stop codons within HvLETM1-like genes (Yao *et al*. 2022; Jayakodi *et al*. 2024). Three LETM1-like proteins were identified: *HORVU.LETM1-like1*, *HORVU.LETM1-like2* and *HORVU.LETM1-like3* (see Fig. S3).

*HORVU.LETM-like1* and *HORVU.LETM1-like2* were found to be expressed uniformly over most tissues, similar to LETM1 domain-containing genes in other plant species, while *HORVU.LETM1-like3* had low to negligible expression (Mascher *et al*. 2017). In the landrace FT880, a four-nucleotide deletion in *HORVU.LETM1-like2* was observed and confirmed. This results in a frameshift at lysine 542 and a subsequent premature stop codon after leucine 568. The mutation is predicted to eliminate the EF-hand domain (Fig. S18). Taf-induced biostimulation was then assessed in FT880 and compared with three control lines (Fig. S19-S20). FT880 exhibited negative responses to Taf treatment, including significant reductions in total, shoot, and root dry weight, whereas the control cultivar Barke exhibited significant increases in root dry weight and a reduced shoot-to-root ratio following Taf inoculation. The control cultivars Bonus and Morex were intermediate. These results support the hypothesis that LETM1-like genes play a key role in mediating Taf-induced biostimulation, and that disruption of these genes can inhibit the beneficial effects of *Trichoderma* inoculation on plant growth.

To directly and quantitatively compare the phenotypic activity of the I/G-type and F-type alleles of *BvCIRA15A/LETM1*, we performed complementation tests in *Atcira15* mutants using the sugar beet alleles from inbred line I and F. We synthesized the coding sequences from the *BvCIRA15A/LETM1.I/G* and *BvCIRA15A/LETM1.F* alleles and the native *AtCIRA15A/LETM1* and cloned into a rescue construct containing the native *AtCIRA15A/LETM1* promoter, 3’ and 5’ UTR, and a NOS termination sequence (Fig. S4-S6). These constructs were then transformed into Col-0, *cira15a-1*, *cira15b-1*, and *cira15b-2* backgrounds. For each case, three independent transformed lines were assessed quantitatively for CIRA responses by electrolyte leakage (Fig. 6A). None of the three constructs interfered with native CIRA function as the transformed Col-0 background performed similarly to previous results and to the untransformed control (Fig. 1B and Fig. 6A). The constructs with the native *AtCIRA15A/LETM1* and the *BvCIRA15A/LETM1*.*I/G* allele were able to fully rescue the CIRA function in *cira15a-1,* demonstrating that the encoded homologs are functionally equivalent. Surprisingly, *cira15b-1* and *cira15b-2* were also rescued by both constructs, demonstrating that *AtCIRA15A/LETM1* and *BvCIRA15A/LETM1*.*I/G* alleles have similar functions to *AtCIRA15B/LETM2* regarding the CIRA response (Fig. 6A). In contrast, the *BvCIRA15A/LETM1.F* allele construct displayed only a partial rescue of the phenotype in all mutants. This response clearly indicates that the *BvCIRA15A/LETM1.F* allele underperforms in allowing CIRA and is the likely cause of the Taf-suppressed growth in sugar beet seedlings in the inbred line F.

**Figure 6.**
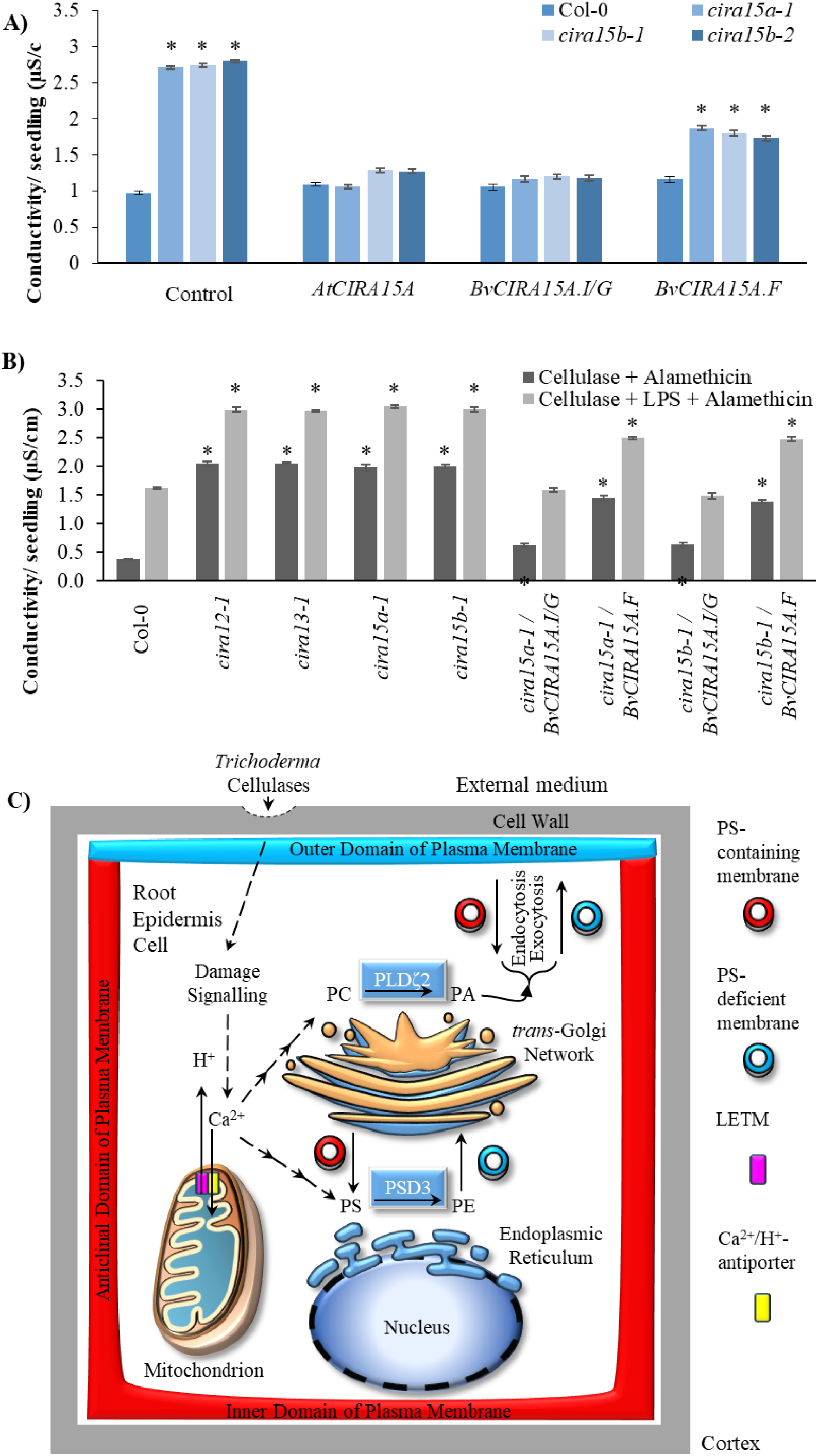
Complementation of *cira15* mutants and lipid-dependent modulation of the CIRA response. (A) The *cira15* mutants display elevated conductivity relative to Col-0, indicating loss of the CIRA response. Expression of *AtCIRA15A* or the sugar beet allele *BvCIRA15A.I/G* restores resistance, whereas *BvCIRA15A.F* provides only partial complementation. Bars represent mean ± SE. Asterisks indicate significant differences compared to Col-0 (FDR q ≤ 0.05; n = 3 biological replicates). (B) Electrolyte leakage in Col-0, lipid metabolism mutants (*cira12-1* and *cira13-1*), *cira15* mutants, and complementation lines expressing *BvCIRA15A* alleles following treatment with cellulase and alamethicin, or cellulase with lysophosphatidylserine (LPS) followed by alamethicin. LPS enhances membrane permeabilization across genotypes while preserving allele-dependent differences. Bars represent mean ± SE. Asterisks indicate significant differences compared to Col-0 within each treatment (FDR q ≤ 0.05; n = 3 biological replicates). (C) Proposed model for LETM involvement in CIRA-associated membrane remodeling and polarity establishment in root epidermal cells. Cellulase-induced damage signaling alters cytosolic Ca²⁺ dynamics, potentially mediated by mitochondrial ion homeostasis. Changes in Ca²⁺ are linked to activation of CIRA13/PLDζ2-dependent vesicular trafficking and CIRA12/PSD3-mediated conversion of phosphatidylserine (PS) to phosphatidylethanolamine (PE) in the endoplasmic reticulum. Vesicles depleted in PS are trafficked to the plasma membrane, contributing to membrane remodeling and resistance to alamethicin. This model is consistent with proposed roles of LETM proteins in Ca²⁺/H⁺ exchange, although their precise function in plants remains unresolved.

To test whether *CIRA15* functions within the broader plasma membrane lipid remodeling processes recently shown to underlie CIRA, we compared *cira15* mutants and complementation lines of *BvCIRA15A.I/G* and *BvCIRA15A.F* with the phospholipid pathway mutants *cira12-1* and *cira13-1* (Panthapulakkal Narayanan *et al*. 2026). Conductivity assays performed after pretreatment with lysophosphatidylserine (LPS), a lysophospholipid previously shown to counteract CIRA by restoring anionic phospholipids in the plasma membrane and altering membrane lipid composition (Panthapulakkal Narayanan *et al*. 2026). The assay revealed that LPS enhanced membrane permeabilization across all *cira12*, *cira13* and *cira15* genotypes while preserving the allele-dependent differences observed for the *BvCIRA15A* alleles (Fig. 6B). These observations indicate that CIRA15 activity is connected to the broader membrane remodeling processes required for CIRA, potentially linking mitochondrial ion homeostasis to lipid-dependent changes in plasma membrane composition (Fig. 6C).

## Discussion

The gene *LETM1* was originally identified as a gene that is deleted in patients with Wolf-Hirschhorn Syndrome (WHS), a human disease involving deficits in multiple developmental and physiological functions, intellectual disability, epilepsy, growth delay, craniofacial dysgenesis, and recently mutations in *HsLETM1* have also been associated with diabetes and several forms of cancer (Endele, Fuhry, Pak, Zabel & Winterpacht 1999; Jiang *et al*. 2009; Li *et al*. 2019; Liu *et al*. 2019; Garbincius & Elrod 2022; Tran *et al*. 2022). Bi-allelic missense and loss-of-function LETM1 variants have further been associated with a complex set of phenotypes reminiscent of mitochondrial diseases, most commonly respiratory chain complex deficiencies, delayed development, and neurological disorders (Kaiyrzhanov *et al*. 2022).

In the light of the medical importance and the complexity of phenotypical syndromes, *LETM1* homologs in especially humans and yeast have been studied biochemically in multiple investigations (Natarajan *et al*. 2021). In humans, the LETM1 function has been suggested to be a Ca^2+^/H^+^ antiporter in the inner membrane of mitochondria maintaining mitochondrial ion homeostasis important for respiration (Austin & Nowikovsky 2019). The assignment of HsLETM1 as a Ca^2+^/H^+^ antiporter has been challenged by the assignment of mammalian TMBIM5 as a Ca^2+^/H^+^ antiporter, indicating that LETM1 may instead be an interaction partner involved in Ca^2+^ and/or K^+^ homeostasis (Austin *et al*. 2022). Though a large number of investigations have not clarified LETM1 function, it has been clearly linked to mitochondrial ion transport and may thus also affect cytosolic Ca^2+^ and/or K^+^ levels (Panda, Behera, Alam & Syed 2021; Garbincius & Elrod 2022). By analogy, it is possible that mitochondrial or cytosolic ion homeostasis effects mediated by plant LETM homologs may influence the CIRA12 and CIRA13 functions that execute the plasma membrane lipid remodeling processes that lead to resistance to alamethicin (Panthapulakkal Narayanan *et al*. 2026).

Outside mammals, phenotypical changes in response to modified LETM1 ortholog expression are consistent with the changes observed in humans. RNAi suppression of the LETM1 ortholog in *Drosophila melanogaster* leads to neurological and mitochondrial dysfunctionality and developmental lethality (McQuibban *et al*. 2010). Likewise, the *Trypanosoma cruzi* LETM1 ortholog is essential for optimal growth, mitochondrial bioenergetics and differentiation (Dos Santos *et al*. 2021; Garbincius & Elrod 2022). Similar to humans, studies in yeast conclude that the LETM1 homolog Mdm38 plays a role central to mitochondrial homeostasis, however yeast homologs prefer K^+^ cations as opposed to Ca^2+^ in humans (Li *et al*. 2019). Based on the conservation of EF hand motives between LETM proteins in animals and plants, but not yeast (Fig. S8), and that plant LETM1 and LETM2 clearly deviates from the yeast homologue by not having a K^+^ binding site (Fig. S16), we would expect to see more similarity in mitochondrial and developmental functions of plant LETM proteins to animal LETM than to yeast Mdm38.

In plants, investigations in *Arabidopsis* have shown both LETM1 domain-containing genes to be essential, as removal of both homologs is lethal (Zhang *et al*. 2012). Interestingly, previous investigators found no significant phenotypic difference to wild type in single mutant lines of *AtCIRA15A/LETM1* or *AtCIRA15/LETM2*, noting that they are potentially acting in redundant roles for general growth (Zhang *et al*. 2012). In contrast, the defect in CIRA induction is here observed in *Arabidopsis* if either of the two genes is disrupted (Fig. 1, Fig. S7B). However, the ability of *AtCIRA15A*/*LETM1* expression to complement the CIRA phenotype of *cira15b* alleles (Fig. 6) indicates functional redundancy, and that potentially the gene dosage may be critical for CIRA induction. In contrast, defects in *HsLETM1* cannot be rescued by expression of the shorter *HsLETM2* homologue, a LETMD-like homolog, indicating that the utility of the EF-Hand domain is required for normal LETM1 function in humans (Tamai *et al*. 2008). The *AtCIRA15B/LETM2* gene displayed parent of origin effects regarding seed development (Zhang *et al*. 2012), and the *HsLETM1* gene displays haploinsufficiency in relation to WHS syndrome (Endele *et al*. 1999). Possibly, the haploinsufficiency of the single *HsLETM1* gene may mirror the need for dual functional *AtCIRA15/LETM* genes for the CIRA process in *Arabidopsis*. In *Arabidopsis*, a double homozygous loss of both *AtCIRA15/LETM1* genes was embryo lethal, and in homo/hemizygous mutant lines an impaired seedling growth and mitochondrial protein accumulation, including oxidative phosphorylation components, was observed (Zhang *et al*. 2012). Therefore, our mechanistic understanding of CIRA15/LETM function in the general mitochondrial context of supporting cellular activity such as cation homeostasis and respiration and how it relates to CIRA is, like for the function-phenotypic connections in other species, still unclear.

Cellulase activity can be perceived by plant cells by monitoring damage-associated molecular patterns (DAMPs), cell wall integrity (CWI), or by binding to the cellulases themselves (Dora, Terrett & Sánchez-Rodríguez 2022). Activity of cellulases has been studied *in vitro* and *in vivo* in great detail, as there are three main types of cellulase activity, endo-cellulase, exo-cellulase and β-glucosidase activity that act in synergy to degrade the main component of the plant cell wall bundles (Jeoh, Cardona, Karuna, Mudinoor & Nill 2017). Previous findings in maize have documented plant beneficial responses to cellulases as the cellulases from *Trichoderma harzianum* (*Thph1* and *Thph2*) induce systemic resistance in the plant (Saravanakumar *et al*. 2016). Likewise, the perception of cellulases and the CIRA response has been linked to plasma membrane reorganization, which leads to resistance to the *Trichoderma spp.* antibiotic peptaibol peptides (Aidemark *et al*. 2010; Dotson *et al*. 2018). Recent work has demonstrated that the CIRA response involves specific remodeling of plasma membrane phospholipids. In *Arabidopsis*, the genes *CIRA12* and *CIRA13*, encoding the phosphatidylserine decarboxylase *PSD3* and the phospholipase *PLDζ2*, respectively, were shown to be required for CIRA and to mediate asymmetric redistribution of anionic phospholipids in the plasma membrane (Panthapulakkal Narayanan *et al*. 2026). These lipid modifications were proposed to increase membrane lipid packing and thereby prevent alamethicin pore formation. CIRA is hypothesized to be a co-symbiotic response by the plant to adjust its membrane constitution to the presence of *Trichoderma spp.* cellulases. These are constitutively expressed as part of the environmental sensing biology of *Trichoderma spp.* and likely function within the symbiotic relationship between plants and *Trichoderma spp*. that promote growth and disease tolerance (Amore, Giacobbe & Faraco 2013; Dotson *et al*. 2018; Chen *et al*. 2024).

Plant biostimulation by microbial agents is an enticing agricultural trait as increases to yield are a cornerstone for plant breeding. A meta-analysis of biostimulation for crop species showed that currently there is only about 10% increase in harvest, as there are variations between crops and biostimulants (Li, Van Gerrewey & Geelen 2022). Ours and previous studies conclude that our current crops, during the long history of breeding, have lost adaptations for symbiosis with biostimulating microbial agents to a various extent (Smith & Goodman 1999; Shoresh & Harman 2008; Tucci *et al*. 2011; Schmidt *et al*. 2020; Zilber-Rosenberg & Rosenberg 2021; Brachi *et al*. 2022). Our studies indicate that there can be selection for enhanced growth of 19% on average with *BvCIRA15A/LETM1* differential markers (Fig. 4B). Therefore, with further investigation into plant and microbial factors that participate in symbiotic communication, large agricultural gains can potentially be reaped.

Our study infers that the ability of plants to cohabitate with a symbiont influences the effects of early biostimulation of the plant by the symbiont. Peptaibols produced by *Trichoderma spp.* can impair root growth (Shi *et al*. 2016), but the stimulus by cellulases from *Trichoderma spp*. induces the CIRA response in both *Arabidopsis* and sugar beet, and leads to an innate resistance by the plants, allowing cohabitation (Dotson *et al*. 2018) (Fig. 1B, 3, and 6). When screening for genes that participate in the CIRA process the gene *AtCIRA15A/LETM1* and the paralog *AtCIRA15B/LETM2* were identified and confirmed to each be crucial for the CIRA response (Fig. 1 and Fig. S7). Associated phenotypes in sugar beet identified variable seedling biostimulation of inbred lines, lending the hypothesis that CIRA participated and was a key part of the biostimulation or symbiosis process. The phenotyping of sequenced inbred sugar beet genomes identified markers that associated with biostimulation and the *BvCIRA15A/LETM1* alleles (Fig. 4 and 5, Fig. S10-S12). Protein modeling of the I/G-and F-type *BvCIRA15A/LETM1* variants suggests reduced function of the F allele, consistent with allele determination and expression analysis identifying *BvCIRA15A/LETM1* allelic differences as a factor significantly associated with the phenotypic disparity between inbred lines (Fig. 3, Fig. S2, and S16). The I/G-type allele types of *BvCIRA15A/LETM1* were associated with early biostimulation and this allele was also able to specifically and fully rescue the CIRA phenotypes of *Atcira15a* and *Atcira15b* mutant lines (Fig. 6). In contrast, the gene product of the *BvCIRA15A/LETM1* F-type allele behaved as a partial-complementation allele and a contributing factor to the negative early biostimulation observed in the sugar beet inbred line F. Supporting the significance of LETM1-containing genes for *T. afroharzianum* T22*-*induced biostimulation, we also find that when mutated in a barley cultivar, significant decreases in growth occurs (Fig. S19-20). Therefore, *CIRA15/LETM* and homologs participate in the cooperation between symbiont and plant, and the identification of an essential plant genetic factor for biocontrol paves the way for future breeding endeavors to adjust crop plants for symbiotic traits such as competence for biostimulation and biocontrol.

## Acknowledgments

The author would like to thank those that have helped this study Oscar Rollano Peñaloza, Johan Pettersson, Sanjana Holla, Ionnis Eilard, Mikko Luomaranta, Egle Kybartaité, Moa Metz, Jesper Persson, Anton Persson, Stefanus Francios Du Trot, and Nathalie Friberg. This research was supported by grants from Carl Tryggers Foundation (AGR), Sven and Lilly Lawski’s fund (SPN), FORMAS research council, The Swedish research council and SLU Grogrund funding.

## Availability of data and materials

Raw sequencing data for the genomes of lines A, B, D, F, G, H and I have been deposited in the European Nucleotide Archive under accession number PRJEB110473. Sequencing data for marker defined regions for all lines are available by request. The datasets used in the current study are available from the corresponding author on reasonable request.

## Authors’ contributions

AGR, BRD, LGB, and KF conceived the investigation and designed the research with expert support from TK, JS, and TE. BRD, MS, SPN and SK performed the experiments. BRD, AGR, SPN, SK, MS, KF, TK, and SB analysed the data with feedback from all other authors. BRD, LGB, and AGR wrote and edited the manuscript, and all authors read and approved the final version of the manuscript.

