## Supplementary material for "Plant LETM1 homologs are required for fungus-induced antibiotic resistance and biostimulation": Supplementary Materials.pdf

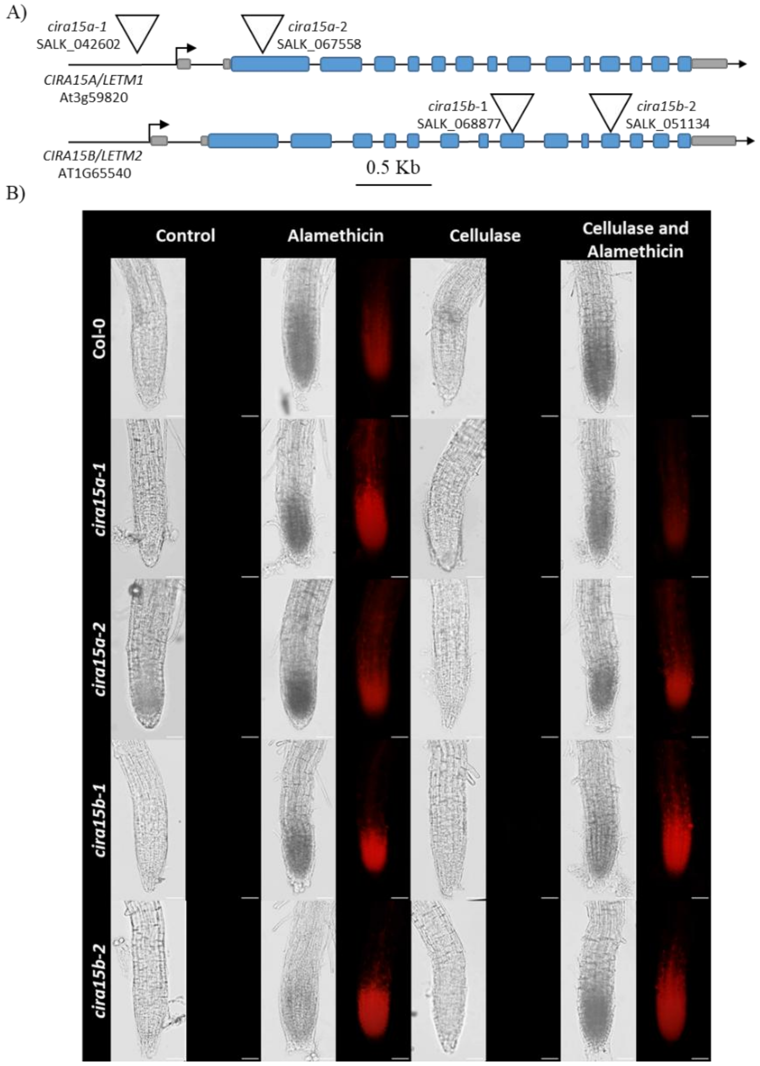


**Supplemental Figure 1: T-DNA mutants of *AtCIRA15A/LETM1A* and *AtCIRA15B/LETM1B* CIRA.** A) Gene model of *AtCIRA15A*/*LETM1* and *AtCIRA15B*/*LETM2* with transcriptional start site (arrow), exon (blue), UTRs (gray), and T-DNA mutations (triangles) modeled. All T-DNA insertion points were verified by PCR using the left border primer and an upstream and downstream native primer. Bar represents 0.5 Kb. B) Representative *Arabidopsis* seedling roots from Col-0, *Atcira15a/letm1a* and *Atcira15b/letm1b* lines treated with cellulase to induced resistance to alamethicin. Three to five-day old seedlings from each line were subjected to treatment control or cellulase (2 hour in 1% *Trichoderma spp.* cellulase) followed by control or 10-minute 20 µg/mL alamethicin treatment. All were then stained with 1.5 µM fluorescent DNA probe propidium iodide (PI) for 1 minute. Both bright field and fluorescent images were captured. Positive staining indicates penetration of Alamethicin into cell membranes and the formation of open channels allowing for the intercalation of PI at double stranded nucleic acids inside the cell. Wild type line (Col-0) demonstrates clear cellulase-induced resistance against alamethicin pore formation whereas all mutant lines of *Atcira15/letm1* remained susceptible upon cellulase treatment. Additional homo/hemi mutant populations of *cira15a/letm1* and *Atcira15b/letm2* were tested and also found to be CIRA deficient. Representative images of 3 independent CIRA assays from pools of at least 3 seedlings. Bars represents 50 µm

| Kingdom | Gene Grouping | Gene count | Species count | Genes per Species | Transmembrane Helix | LETM/MDM38-like | LETM1 RBD | N terminal Coiled Coils | C Terminal Coiled Coils | EF-Hand domains | EF-Hand Pair |
|---|---|---|---|---|---|---|---|---|---|---|---|
| Plantae | LETM1 EF-Hand - like | 59 | 35 | 1.7 | 1.0 | 1.0 | 1.0 | 0.0 | 2.3 | 1.1 | 1.0 |
| Animalia | LETM1 EF-Hand - like | 6 | 6 | 1.0 | 1.0 | 1.0 | 1.0 | 0.5 | 3.7 | 1.2 | 0.8 |
| Fungi | LETM1 EF-Hand - like | 2 | 6 | 0.3 | 1.0 | 1.0 | 1.0 | 0.0 | 3.0 | 0.5 | 1.0 |
| Heterokont | LETM1 EF-Hand - like | 1 | 1 | 1.0 | 1.0 | 1.0 | 1.0 | 1.0 | 2.0 | 2.0 | 1.0 |
| Plantae | LETMD - like | 9 | 35 | 0.3 | 0.6 | 1.1 | 0.7 | 0.1 | 0.4 | - | - |
| Animalia | LETMD - like | 10 | 6 | 1.7 | 0.9 | 1.0 | 1.0 | 0.2 | 0.1 | - | - |
| Fungi | LETMD - like | 5 | 6 | 0.8 | 1.0 | 1.0 | 1.0 | 0.6 | 1.6 | - | - |
| Heterokont | LETMD - like | 0 | 1 | - | - | - | - | - | - | - | - |

**Supplemental Figure 2: Summary of gene components of LETM1 domain-containing proteins across kingdoms.** Count genes represents the number of genes identified in the kingdom. Count species represents the number of species surveyed in the kingdom. Genes/Species represents the average number of genes per species. Other columns are average number of respective domains per gene within each group. All protein sequences were analyzed within the program Geneious (Kearse *et al.* 2012) predicted domains predicted are Transmembrane Helix denoted by TMHMM, LETM1/MDM38-like (IPR044202), LETM1 RBD (IPR033122), N and C terminal Coiled Coils denoted by Coils, EF-Hand domain (IPR002048) and EF-Hand Pair (IPR011992). Cells are colored according to relative abundance. For complete list of LETM homologs see Fig. S4.

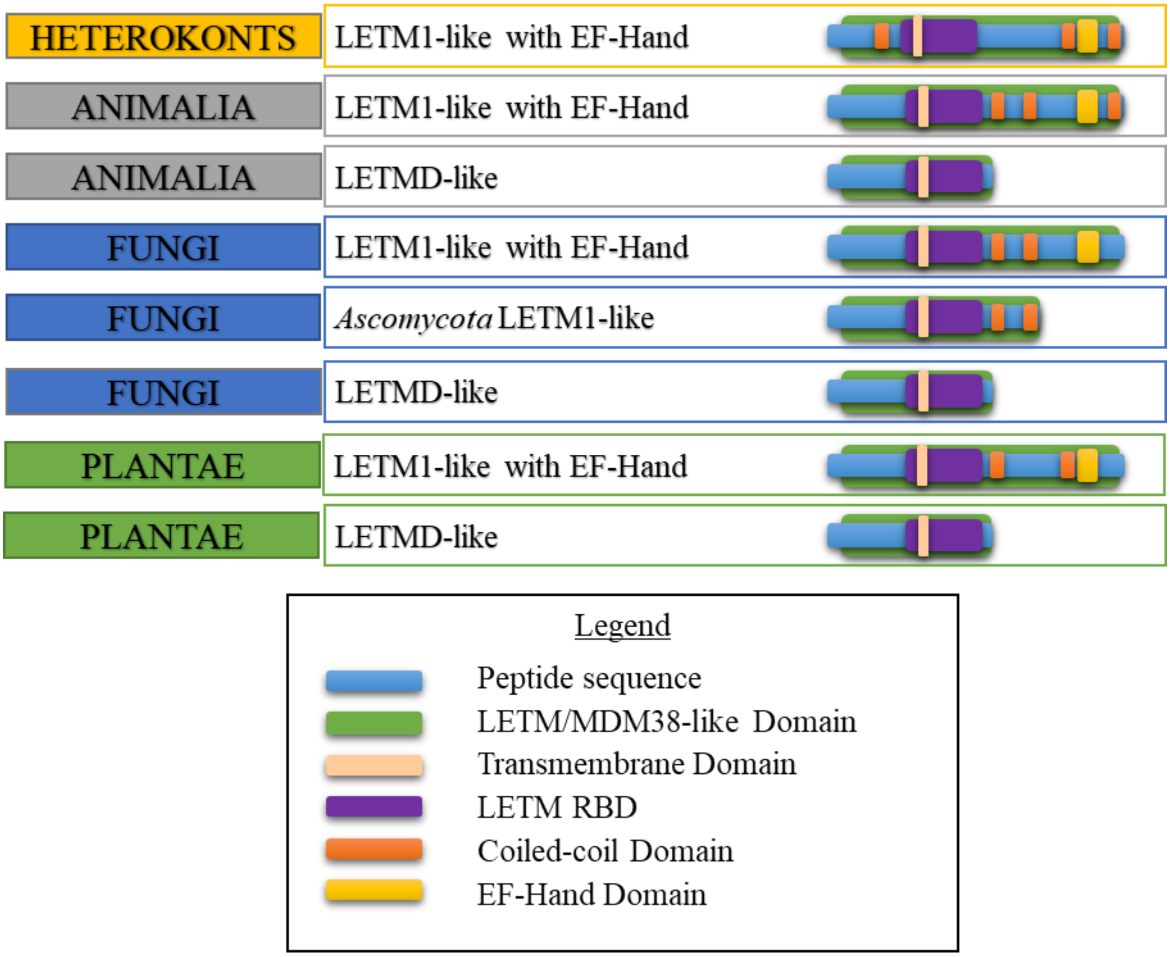


**Supplemental Figure 3: Domain analysis of LETM1 domain-containing genes in Eukaryotes.** Graphical representation of different groupings of LETM1 domain genes from heterokonts, animalia, fungi and plantae kingdoms. Apart from some fungal clades, all genomes contained at least one of the LETM1-like and EF Hand-domain genes that also contain LETM1 ribosomal binding domain, at least one transmembrane domain, and variable amounts of coiled coil domains. Some fungal species, particularly in *Ascomycota*, lacked an identifiable EF-Hand domain. Additionally, several species of plants and fungi contain genes encoding proteins with a reduced LETM1 signature, lacking coiled coil and EF-Hand domains. These are similar to human *HsLETM2* and *HsLETMD* genes and therefore are termed LETMD-like here. For complete list of LETM homologs see Fig. S4.

| Abreviated gene name | Protein ID | Organism | Kingdom | Phylum-Order |
|---|---|---|---|---|
| PHYPAR.LETM1-like1 | W2QWZ9 | *Phytophthora parasitica* | Heterokont | Oomycota |
| DROME.LETM1-like1 | P91927 | *Drosophila melanogaster* | Animalia | Arthropoda |
| CAEEL.LETM1-like1 | Q9XVM0 | *Caenorhabditis elegans* | Animalia | Nemotoda |
| CAEEL.LETMD-like1 | Q8I4K3 | *Caenorhabditis elegans* | Animalia | Nemotoda |
| DROME.LETMD-like1 | Q9VSM4 | *Drosophila melanogaster* | Animalia | Arthropoda |
| STRPU.LETM1-like1 | A0A7M7NWP2 | *Strongylocentrotus purpuratus* | Animalia | Echinodermata |
| STRPU.LETMD-like1 | A0A7M7RFP5 | *Strongylocentrotus purpuratus* | Animalia | Echinodermata |
| STRPU.LETMD-like2 | A0A7M7N3J9 | *Strongylocentrotus purpuratus* | Animalia | Echinodermata |
| DANRE_LETM1-like1 | Q1LY46 | *Danio rerio* | Animalia | Chordata |
| DANRE_LETMD-like1 | F1QSQ8 | *Danio rerio* | Animalia | Chordata |
| DANRE_LETMD-like2 | B3DJY5 | *Danio rerio* | Animalia | Chordata |
| MUSMU.LETM1-like1 | Q9Z2I0 | *Mus musculus* | Animalia | Chordata |
| MUSMU.LETMD-like1 | Q7TNU7 | *Mus musculus* | Animalia | Chordata |
| MUSMU.LETMD-like2 | Q924L1 | *Mus musculus* | Animalia | Chordata |
| HOMSA.LETM1 | O95202 | *Homo sapiens* | Animalia | Chordata |
| HOMSA.LETMD-like1 | Q2VYF4 | *Homo sapiens* | Animalia | Chordata |
| HOMSA.LETMD-like2 | Q6P1Q0 | *Homo Sapiens* | Animalia | Chordata |
| BATDE.LETM1-like1 | F4P1E1 | *Batrachochytrium dendrobatidis* | Fungi | Chytridiomycota |
| CRYNE.LETMD-like1 | UOH80244.1 | *Cryptococcus neoformans* | Fungi | Basidiomycota |
| RHIZO.LETM1-like1 | CEL59443.1 | *Rhizoctonia solani* | Fungi | Basidiomycota |
| SACCE.MDM38-LETMD-like1 | Q08179 | *Saccharomyces cerevisiae* | Fungi | Ascomycota |
| SACCE.YLH47.LETMD-like2 | Q06493 | *Saccharomyces cerevisiae* | Fungi | Ascomycota |
| FUSGR.LETMD-like1 | EYB26808.1 | *Fusarium graminearum* | Fungi | Ascomycota |
| TRIHA.LETMD-like1 | A0A2T4ACP3 | *Trichoderma harzianum* | Fungi | Ascomycota |
| PORUM.LETM1-like1 | Pum0513s0001.1 | *Porphyra umbilicalis* | Plantae | Rhodophyta |
| KLENI.LETM1-like1 | A0A1Y1I912 | *Klebsormidium nitens* | Plantae | Charophyta |
| BOTBR.LETM1-like1 | Bobra.0258s0003.1 | *Botryococcus braunii* | Plantae | Chlorophyta |
| CHLRE.LETM1-like1 | A0A2K3CWR6 | *Chlamydomonas reinhardtii* | Plantae | Chlorophyta |
| COCSU.LETM1-like1 | COCSUDRAFT_49147 | *Coccomyxa subellipsoidea* | Plantae | Chlorophyta |
| OSTLU.LETM1-like1 | OSTLU_50194 | *Ostreococcus lucimarinus* | Plantae | Chlorophyta |
| VOLCA.LETM1-like1 | Vocar.0043s0044.1 | *Volvox carteri* | Plantae | Chlorophyta |
| CERPU.LETM1-like1 | CepurGG1.3G126600.1 | *Ceratodon purpureus* | Plantae | Bryophyta |
| PHYPA.LETM1-like1 | Pp3c11_19590V3.1 | *Physcomitrella patens* | Plantae | Bryophyta |
| PHYPA.LETMD-like1 | Pp3c23_15550V3.1 | *Physcomitrium patens* | Plantae | Bryophyta |
| SPHFA.LETM1-like1 | Sphfalx16G020700.1 | *Sphagnum fallax* | Plantae | Bryophyta |
| SPHFA.LETM1-like2 | Sphfalx17G066900.1 | *Sphagnum fallax* | Plantae | Bryophyta |
| SPHMA.LETM1-like1 | Sphmag16G019300.1 | *Sphagnum magellanicum* | Plantae | Bryophyta |
| SPHMA.LETM1-like2 | Sphmag17G065100.1 | *Sphagnum magellanicum* | Plantae | Bryophyta |
| MARPO.LETM1-like1 | Mapoly0198s0012.1 | *Marchantia polymorpha* | Plantae | Bryophyta |
| DIPCO.LETM1-like1 | Dicom.13G083600.2 | *Diphasiastrum complanatum* | Plantae | Lycophytes |
| DIPCO.LETM1-like2 | Dicom.13G085300.1 | *Diphasiastrum complanatum* | Plantae | Lycophytes |
| DIPCO.LETM1-like3 | Dicom.03G115700.2 | *Diphasiastrum complanatum* | Plantae | Lycophytes |
| SELMO.LETM1-like1 | D8RVZ0 | *Selaginella moellendorffii* | Plantae | Lycophytes |
| CERRI.LETM1-like1 | Ceric.01G129300.1 | *Ceratopteris richardii* | Plantae | Polypodiophyta |
| CERRI.LETM1-like2 | Ceric.23G078200.1 | *Ceratopteris richardii* | Plantae | Polypodiophyta |
| CERRI.LETMD-like1 | Ceric.35G053400.1 | *Ceratopteris richardii* | Plantae | Polypodiophyta |
| THUPL.LETM1-like1 | Thupl.29382393s0001.1 | *Thuja plicata* | Plantae | Gymnosperms |
| THUPL.LETMD-like1 | Thupl.29380438s0004.1 | *Thuja plicata* | Plantae | Gymnosperms |
| NYMCO.LETM1-like1 | Nycol.N00027.1 | *Nymphaea colorata* | Plantae | Angiosperms-Nymphaeales |
| AMBTR.LETM1-like1 | AMTR_s00066p00145760 | *Amborella trichopoda* | Plantae | Angiosperms-Amborellales |
| CINKA.LETM1-like1 | CKAN_02614300 | *Cinnamomum kanehirae* | Plantae | Angiosperms -Laurales |
| CINKA.LETM1-like2 | CKAN_01270100 | *Cinnamomum kanehirae* | Plantae | Angiosperms -Laurales |
| CINKA.LETMD-like1 | CKAN_00175900 | *Cinnamomum kanehirae* | Plantae | Angiosperms -Laurales |
| ASPCO.LETM1-like1 | evm.model.AsparagusV1_09.515 | *Asparagus officinalis* | Plantae | Angiosperms-Monocot |
| ASPCO.LETM1-like2 | evm.model.AsparagusV1_02.498 | *Asparagus officinalis* | Plantae | Angiosperms-Monocot |
| ASPCO.LETMD-like1 | evm.model.AsparagusV1_07.2805 | *Asparagus officinalis* | Plantae | Angiosperms-Monocot |
| ASPCO.LETMD-like2 | evm.model.AsparagusV1_10.316 | *Asparagus officinalis* | Plantae | Angiosperms-Monocot |
| BRADI.LETM1-like1 | Bradi2g26300.1. | *Brachypodium distachyon* | Plantae | Angiosperms-Monocot |
| BRADI.LETM1-like2 | Bradi3g48350.2 | *Brachypodium distachyon* | Plantae | Angiosperms-Monocot |
| BRADI.LETM1-like3 | Bradi5g14570.1 | *Brachypodium distachyon* | Plantae | Angiosperms-Monocot |
| HORVU.LETM1-like1 | HORVU.MOREX.r3.1HG0064010.1 | *Hordeum vulgare subsp. vulgare* | Plantae | Angiosperms-Monocot |
| HORVU.LETM1-like2 | HORVU.MOREX.r3.6HG0593260.1 | *Hordeum vulgare subsp. vulgare* | Plantae | Angiosperms-Monocot |
| HORVU.LETM1-like3 | HORVU.MOREX.r3.2HG0178460.1 | *Hordeum vulgare subsp. vulgare* | Plantae | Angiosperms-Monocot |
| MUSCO.LETM1-like1 | A0A804INR5 | *Musa acuminata subsp. malaccensis* | Plantae | Angiosperms-Monocot |
| ORYSA.LETM1-like1 | Q6K1Y0 | *Oryza sativa subsp. japonica* | Plantae | Angiosperms-Monocot |
| ORYSA.LETM1-like2 | Q7XUK2 | *Oryza sativa subsp. japonica* | Plantae | Angiosperms-Monocot |
| AQUCO.LETM1-like1 | AQUCO_01600234v1 | *Aquilegia coerulea* | Plantae | Angiosperms-Eudicot |
| ARATH.CIRA15A.LET1A | Q9M1Z2 | *Arabidopsis thaliana* | Plantae | Angiosperms-Eudicot |
| ARATH.CIRA15B.LETM1B | F4IBH5 | *Arabidopsis thaliana* | Plantae | Angiosperms-Eudicot |
| BETVU.CIRA15A.LETM1A | KMT15550.1 | *Beta vulgaris subsp. vulgaris* | Plantae | Angiosperms-Eudicot |
| BETVU.CIRA15B.LET1MB | KMT14692 | *Beta vulgaris subsp. vulgaris* | Plantae | Angiosperms-Eudicot |
| CHEQU.LETM1-like1 | LOC110701043 | *Chenopodium quinoa* | Plantae | Angiosperms-Eudicot |
| CHEQU.LETM1-like2 | LOC110718273 | *Chenopodium quinoa* | Plantae | Angiosperms-Eudicot |
| CHEQU.LETM1-like3 | LOC110736374 | *Chenopodium quinoa* | Plantae | Angiosperms-Eudicot |
| CHEQU.LETM1-like4 | LOC110738681 | *Chenopodium quinoa* | Plantae | Angiosperms-Eudicot |
| LOTJA.LETM1-like1 | Lj1g0025516.1 | *Lotus japonicus* | Plantae | Angiosperms-Eudicot |
| LOTJA.LETM1-like2 | Lj4g0005716.1 | *Lotus japonicus* | Plantae | Angiosperms-Eudicot |
| MEDTR.LETM1-like1 | G7KSF6 | *Medicago truncatula* | Plantae | Angiosperms-Eudicot |
| MEDTR.LETM1-like2 | G7L8S9 | *Medicago truncatula* | Plantae | Angiosperms-Eudicot |
| MIMGU.LETM1-like1 | MIMGU_mgv1a001786mg | *Mimulus guttatus* | Plantae | Angiosperms-Eudicot |
| MIMGU.LETM1-like2 | MIMGU_mgv1a001769mg | *Mimulus guttatus* | Plantae | Angiosperms-Eudicot |
| PONTR.LETM1-like1 | Ptrif.0003s2494.2 | *Poncirus trifoliata* | Plantae | Angiosperms-Eudicot |
| PONTR.LETM1-like2 | Ptrif.0002s2197.1 | *Poncirus trifoliata* | Plantae | Angiosperms-Eudicot |
| POPTR.LETM1-like1 | Potri.008G078600.1 | *Populus trichocarpa* | Plantae | Angiosperms-Eudicot |
| POPTR.LETM1-like2 | Potri.007G142000.1 | *Populus trichocarpa* | Plantae | Angiosperms-Eudicot |
| POPTR.LETMD-like1 | Potri.010G178400.1 | *Populus trichocarpa* | Plantae | Angiosperms-Eudicot |
| SOLLY.LETM1-like1 | Solyc12g096810.2.1 | *Solanum lycopersicum* | Plantae | Angiosperms-Eudicot |
| SOLLY.LETM1-like2 | Solyc11g008770.2.1 | *Solanum lycopersicum* | Plantae | Angiosperms-Eudicot |
| VITVI.LETM1-like1 | VIT_205s0094g01070.2 | *Vitis vinifera* | Plantae | Angiosperms-Eudicot |
| VITVI.LETMD-like1 | VIT_200s0220g00090.1 | *Vitis vinifera* | Plantae | Angiosperms-Eudicot |
| VITVI.LETMD-like2 | VIT_209s0002g03320.1 | *Vitis vinifera* | Plantae | Angiosperms-Eudicot |
| VITVI.LETMD-like3 | VIT_209s0002g03310.1 | *Vitis vinifera* | Plantae | Angiosperms-Eudicot |

**Supplemental Figure 4: Table of genes used in the study.** Associated accession abbreviated names, protein ID, organisms, kingdoms and phylum/order of genes used in this study.

| Inbred line | Treatment | N | Total Weight Log2 Norm to Control | Shoot Weight Log2 Norm to Control | Root Weight Log2 Norm to Control | Primary Root Length Log2 Norm to Control | Root Weight per Length Log2 Norm to Control | Shoot to Root Ratio Log2 Norm to Control | Marker - IIS3758 | Marker - IIS8436 |
|---|---|---|---|---|---|---|---|---|---|---|
| I | C | 24 | 0.00 ± 0.07 | 0.00 ± 0.07 | 0.00 ± 0.07 | 0.00 ± 0.05 | 0.00 ± 0.12 | 0.00 ± 0.05 | A/C | G/G |
| I | Taf | 19 | 0.47 ± 0.07 | 0.33 ± 0.06 | 0.67 ± 0.16 | 0.05 ± 0.07 | 0.58 ± 0.1 | -0.11 ± 0.11 | | |
| 55 | C | 12 | 0.00 ± 0.12 | 0.00 ± 0.08 | 0.00 ± 0.2 | 0.00 ± 0.07 | 0.00 ± 0.16 | 0.00 ± 0.13 | _/_ | G/G |
| 55 | Taf | 12 | 0.03 ± 0.13 | 0.13 ± 0.09 | 0.23 ± 0.36 | -0.06 ± 0.07 | 0.14 ± 0.35 | 0.41 ± 0.28 | | |
| 57 | C | 16 | 0.00 ± 0.04 | 0.00 ± 0.05 | 0.00 ± 0.08 | 0.00 ± 0.12 | 0.00 ± 0.16 | 0.00 ± 0.07 | _/_ | G/G |
| 57 | Taf | 12 | 0.29 ± 0.05 | 0.22 ± 0.09 | 0.07 ± 0.06 | -0.45 ± 0.18 | 0.62 ± 0.21 | 0.25 ± 0.13 | | |
| 59 | C | 12 | 0.00 ± 0.13 | 0.00 ± 0.15 | 0.00 ± 0.09 | 0.00 ± 0.05 | 0.00 ± 0.05 | 0.00 ± 0.09 | _/_ | G/G |
| 59 | Taf | 12 | 0.3 ± 0.06 | 0.27 ± 0.06 | 0.27 ± 0.12 | 0.2 ± 0.09 | 0.27 ± 0.08 | -0.09 ± 0.15 | | |
| 66 | C | 13 | 0.00 ± 0.07 | 0.00 ± 0.08 | 0.00 ± 0.16 | 0.00 ± 0.05 | 0.00 ± 0.14 | 0.00 ± 0.15 | C/C | G/G |
| 66 | Taf | 14 | 0.47 ± 0.09 | 0.12 ± 0.07 | 0.5 ± 0.19 | 0.22 ± 0.17 | -0.14 ± 0.22 | -0.29 ± 0.17 | | |
| 71 | C | 12 | 0.00 ± 0.11 | 0.00 ± 0.12 | 0.00 ± 0.12 | 0.00 ± 0.05 | 0.00 ± 0.07 | 0.00 ± 0.1 | _/_ | G/G |
| 71 | Taf | 12 | 0.14 ± 0.07 | 0.26 ± 0.11 | 0 ± 0.09 | -0.09 ± 0.12 | -0.26 ± 0.16 | 0 ± 0.08 | | |
| G | C | 28 | 0.00 ± 0.06 | 0.00 ± 0.06 | 0.00 ± 0.05 | 0.00 ± 0.06 | 0.00 ± 0.07 | 0.00 ± 0.07 | C/C | G/G |
| G | Taf | 24 | 0.45 ± 0.11 | 0.52 ± 0.1 | 0.58 ± 0.16 | -0.16 ± 0.11 | 0.8 ± 0.1 | -0.31 ± 0.08 | | |
| 49 | C | 16 | 0.00 ± 0.07 | 0.00 ± 0.07 | 0.00 ± 0.1 | 0.00 ± 0.05 | 0.00 ± 0.05 | 0.00 ± 0.06 | _/_ | G/G |
| 49 | Taf | 16 | -0.02 ± 0.08 | -0.03 ± 0.1 | 0.04 ± 0.1 | 0.24 ± 0.1 | -0.2 ± 0.07 | -0.15 ± 0.1 | | |
| 60 | C | 18 | 0.00 ± 0.08 | 0.00 ± 0.1 | 0.00 ± 0.05 | 0.00 ± 0.06 | 0.00 ± 0.03 | 0.00 ± 0.09 | C/C | G/G |
| 60 | Taf | 18 | 0.21 ± 0.04 | 0.1 ± 0.04 | 0.34 ± 0.06 | -0.11 ± 0.09 | 0.41 ± 0.1 | -0.14 ± 0.08 | | |
| 78 | C | 14 | 0.00 ± 0.08 | 0.00 ± 0.08 | 0.00 ± 0.09 | 0.00 ± 0.05 | 0.00 ± 0.12 | 0.00 ± 0.17 | _/_ | G/G |
| 78 | Taf | 14 | 0.11 ± 0.13 | 0.33 ± 0.13 | 0.09 ± 0.13 | 0.35 ± 0.12 | 0.15 ± 0.13 | 0.43 ± 0.11 | | |
| A | C | 28 | 0.00 ± 0.07 | 0.00 ± 0.07 | 0.00 ± 0.09 | 0.00 ± 0.04 | 0.00 ± 0.07 | 0.00 ± 0.04 | A/C | G/G |
| A | Taf | 20 | 0.3 ± 0.11 | 0.28 ± 0.09 | 0.08 ± 0.11 | 0.07 ± 0.05 | -0.03 ± 0.11 | -0.14 ± 0.06 | | |
| 52 | C | 16 | 0.00 ± 0.06 | 0.00 ± 0.07 | 0.00 ± 0.06 | 0.00 ± 0.07 | 0.00 ± 0.05 | 0.00 ± 0.09 | _/_ | G/G |
| 52 | Taf | 16 | -0.05 ± 0.05 | -0.05 ± 0.06 | -0.15 ± 0.1 | -0.07 ± 0.09 | -0.02 ± 0.07 | 0.16 ± 0.11 | | |
| 58 | C | 19 | 0.00 ± 0.04 | 0.00 ± 0.04 | 0.00 ± 0.09 | 0.00 ± 0.04 | 0.00 ± 0.07 | 0.00 ± 0.09 | _/_ | G/G |
| 58 | Taf | 18 | -0.18 ± 0.06 | -0.21 ± 0.07 | -0.11 ± 0.12 | -0.1 ± 0.09 | 0.17 ± 0.14 | -0.12 ± 0.12 | | |
| H | C | 21 | 0.00 ± 0.07 | 0.00 ± 0.08 | 0.00 ± 0.07 | 0.00 ± 0.06 | 0.00 ± 0.08 | 0.00 ± 0.06 | A/C | G/G |
| H | Taf | 15 | 0.28 ± 0.1 | 0.17 ± 0.12 | 0.57 ± 0.11 | -0.04 ± 0.13 | 0.6 ± 0.1 | -0.34 ± 0.12 | | |
| 46 | C | 16 | 0.00 ± 0.07 | 0.00 ± 0.03 | 0.00 ± 0.11 | 0.00 ± 0.05 | 0.00 ± 0.11 | 0.00 ± 0.05 | _/_ | G/G |
| 46 | Taf | 15 | 0.24 ± 0.04 | 0.22 ± 0.05 | 0.2 ± 0.08 | 0.06 ± 0.05 | 0.32 ± 0.11 | -0.16 ± 0.09 | | |
| 51 | C | 18 | 0.00 ± 0.06 | 0.00 ± 0.05 | 0.00 ± 0.08 | 0.00 ± 0.04 | 0.00 ± 0.08 | 0.00 ± 0.04 | C/C | G/G |
| 51 | Taf | 16 | 0.04 ± 0.06 | 0.12 ± 0.06 | 0.01 ± 0.09 | 0.13 ± 0.08 | -0.11 ± 0.07 | 0.22 ± 0.04 | | |
| 65 | C | 17 | 0.00 ± 0.12 | 0.00 ± 0.11 | 0.00 ± 0.21 | 0.00 ± 0.06 | 0.00 ± 0.15 | 0.00 ± 0.15 | A/A | G/G |
| 65 | Taf | 16 | 0.75 ± 0.08 | 0.69 ± 0.08 | 0.42 ± 0.06 | 0.14 ± 0.06 | 0.31 ± 0.07 | -0.17 ± 0.09 | | |
| F | C | 20 | 0.00 ± 0.08 | 0.00 ± 0.06 | 0.00 ± 0.12 | 0.00 ± 0.05 | 0.00 ± 0.1 | 0.00 ± 0.07 | C/C | A/A |
| F | Taf | 16 | -0.35 ± 0.05 | -0.33 ± 0.08 | -0.32 ± 0.1 | -0.45 ± 0.11 | 0.16 ± 0.12 | -0.13 ± 0.11 | | |
| 50 | C | 16 | 0.00 ± 0.05 | 0.00 ± 0.05 | 0.00 ± 0.05 | 0.00 ± 0.06 | 0.00 ± 0.06 | 0.00 ± 0.06 | C/C | A/A |
| 50 | Taf | 15 | -0.22 ± 0.05 | -0.11 ± 0.04 | -0.15 ± 0.13 | -0.06 ± 0.08 | -0.23 ± 0.13 | 0.07 ± 0.11 | | |
| 54 | C | 12 | 0.00 ± 0.13 | 0.00 ± 0.1 | 0.00 ± 0.13 | 0.00 ± 0.09 | 0.00 ± 0.16 | 0.00 ± 0.24 | _/_ | A/A |
| 54 | Taf | 12 | 0.22 ± 0.14 | 0.23 ± 0.07 | -0.25 ± 0.15 | 0.13 ± 0.05 | -0.23 ± 0.15 | -0.07 ± 0.14 | | |
| 56 | C | 12 | 0.00 ± 0.02 | 0.00 ± 0.05 | 0.00 ± 0.09 | 0.00 ± 0.1 | 0.00 ± 0.08 | 0.00 ± 0.1 | _/_ | A/A |
| 56 | Taf | 12 | -0.1 ± 0.09 | -0.15 ± 0.06 | -0.4 ± 0.19 | 0.18 ± 0.07 | -0.71 ± 0.18 | 0.45 ± 0.14 | | |
| 77 | C | 14 | 0.00 ± 0.07 | 0.00 ± 0.1 | 0.00 ± 0.11 | 0.00 ± 0.1 | 0.00 ± 0.09 | 0.00 ± 0.13 | C/C | A/A |
| 77 | Taf | 14 | 0.31 ± 0.11 | 0.24 ± 0.12 | 0.14 ± 0.16 | 0.19 ± 0.14 | -0.18 ± 0.17 | -0.18 ± 0.12 | | |
| B | C | 17 | 0.00 ± 0.09 | 0.00 ± 0.09 | 0.00 ± 0.08 | 0.00 ± 0.09 | 0.00 ± 0.1 | 0.00 ± 0.09 | C/T | G/G |
| B | Taf | 12 | -0.18 ± 0.09 | -0.24 ± 0.14 | 0.23 ± 0.08 | 0.19 ± 0.05 | -0.16 ± 0.11 | -0.04 ± 0.06 | | |
| D | C | 30 | 0.00 ± 0.05 | 0.00 ± 0.06 | 0.00 ± 0.06 | 0.00 ± 0.03 | 0.00 ± 0.1 | 0.00 ± 0.06 | A/A | G/G |
| D | Taf | 24 | 0 ± 0.04 | 0 ± 0.06 | 0.26 ± 0.09 | -0.24 ± 0.08 | 0.41 ± 0.1 | -0.33 ± 0.1 | | |
| 63 | C | 13 | 0.00 ± 0.1 | 0.00 ± 0.13 | 0.00 ± 0.17 | 0.00 ± 0.11 | 0.00 ± 0.04 | 0.00 ± 0.22 | _/_ | G/G |
| 63 | Taf | 13 | -0.36 ± 0.09 | -0.22 ± 0.04 | 0.05 ± 0.29 | 0.13 ± 0.09 | -0.17 ± 0.23 | -0.31 ± 0.19 | | |
| E | C | 23 | 0.00 ± 0.11 | 0.00 ± 0.09 | 0.00 ± 0.17 | 0.00 ± 0.1 | 0.00 ± 0.06 | 0.00 ± 0.05 | C/C | A/A |
| E | Taf | 24 | 0.12 ± 0.1 | 0.01 ± 0.1 | 0.39 ± 0.13 | 0.18 ± 0.08 | 0.75 ± 0.09 | -0.32 ± 0.09 | | |
| lines with N<12 | | | | | | | | | | |
| 02 | C | 1 | 0.00 ± NA | 0.00 ± NA | 0.00 ± NA | 0.00 ± NA | 0.00 ± NA | 0.00 ± NA | _/_ | A/A |
| 02 | Taf | 3 | -0.49 ± 0.34 | 0.44 ± 0.59 | 0.62 ± 0.21 | 0.06 ± 0.38 | 0.27 ± 0.12 | 0.11 ± 0.09 | | |
| 05 | C | 5 | 0.00 ± 0.14 | 0.00 ± 0.17 | 0.00 ± 0.49 | 0.00 ± 0.31 | 0.00 ± 0.43 | 0.00 ± 0.48 | _/_ | A/A |
| 05 | Taf | 6 | -0.16 ± 0.08 | 0.05 ± 0 | 0.15 ± 0.07 | 0.08 ± 0.21 | 0.68 ± 0.11 | -0.44 ± 0.22 | | |
| 31 | C | 4 | 0.00 ± 0.06 | 0.00 ± 0.02 | 0.00 ± 0.1 | 0.00 ± 0.15 | 0.00 ± 0.01 | 0.00 ± 0.13 | A/A | G/G |
| 31 | Taf | 4 | -0.03 ± 0.1 | -0.01 ± 0.19 | -0.26 ± 0 | -0.21 ± 0.22 | -0.07 ± 0.16 | -0.17 ± 0.35 | | |
| 47 | C | 10 | 0.00 ± 0.2 | 0.00 ± 0.2 | 0.00 ± 0.15 | 0.00 ± 0.09 | 0.00 ± 0.1 | 0.00 ± 0.13 | A/C | G/G |
| 47 | Taf | 10 | 0.26 ± 0.2 | 0.25 ± 0.21 | 0.55 ± 0.17 | -0.03 ± 0.16 | 0.18 ± 0.07 | 0.02 ± 0.06 | | |
| 48 | C | 11 | 0.00 ± 0.1 | 0.00 ± 0.1 | 0.00 ± 0.12 | 0.00 ± 0.11 | 0.00 ± 0.1 | 0.00 ± 0.12 | C/C | A/G |
| 48 | Taf | 14 | 0.28 ± 0.09 | 0.18 ± 0.08 | 0.55 ± 0.12 | 0.29 ± 0.06 | 0.46 ± 0.11 | -0.2 ± 0.09 | | |
| 64 | C | 4 | 0.00 ± 0.14 | 0.00 ± 0.09 | 0.00 ± 0.27 | 0.00 ± 0.03 | 0.00 ± 0.11 | 0.00 ± 0.14 | _/_ | G/G |
| 64 | Taf | 4 | 0.4 ± 0.08 | 0.23 ± 0.19 | 0.75 ± 0.11 | 0.13 ± 0 | 0.8 ± 0.04 | -0.7 ± 0.3 | | |
| 68 | C | 4 | 0.00 ± 0.08 | 0.00 ± 0.01 | 0.00 ± 0.08 | 0.00 ± 0.16 | 0.00 ± 0.01 | 0.00 ± 0.06 | _/_ | G/G |
| 68 | Taf | 4 | 0.36 ± 0.14 | -0.04 ± 0.19 | 0.51 ± 0.07 | 0.44 ± 0.2 | -0.03 ± 0.16 | -0.47 ± 0.04 | | |
| 69 | C | 2 | 0.00 ± 0.36 | 0.00 ± 0.47 | 0.00 ± 0 | 0.00 ± 0.5 | 0.00 ± 0.5 | 0.00 ± 0.47 | C/C | G/G |
| 69 | Taf | 4 | 1.51 ± 0.52 | 1.41 ± 0.36 | 1.6 ± 0.93 | 1.38 ± 0.37 | 0.22 ± 0.58 | -0.19 ± 0.58 | | |
| 70 | C | 6 | 0.00 ± 0.21 | 0.00 ± 0.25 | 0.00 ± 0.37 | 0.00 ± 0.09 | 0.00 ± 0.33 | 0.00 ± 0.33 | _/_ | G/G |
| 70 | Taf | 6 | 0.67 ± 0.33 | 0.57 ± 0.34 | 1.06 ± 0.26 | -0.38 ± 0.17 | 1.41 ± 0.33 | -0.8 ± 0.07 | | |
| 73 | C | 7 | 0.00 ± 0.11 | 0.00 ± 0.13 | 0.00 ± 0.12 | 0.00 ± 0.23 | 0.00 ± 0.27 | 0.00 ± 0.14 | _/_ | G/G |
| 73 | Taf | 7 | -0.08 ± 0.11 | -0.06 ± 0.1 | -0.09 ± 0.2 | 0.32 ± 0.11 | -0.12 ± 0.14 | 0.07 ± 0.21 | | |
| 72 | C | 6 | 0.00 ± 0.26 | 0.00 ± 0.22 | 0.00 ± 0.25 | 0.00 ± 0.13 | 0.00 ± 0.05 | 0.00 ± 0.18 | _/_ | G/G |
| 72 | Taf | 6 | -0.11 ± 0.03 | -0.25 ± 0.08 | -0.28 ± 0.21 | 0.1 ± 0.29 | -0.36 ± 0.06 | 0.47 ± 0.07 | | |

**Supplemental Figure 5: Table of inbred line phenotypes and markers used in the study.** Associated Refbeet1.5 markers and Log2 transformed and normalized to control phenotypes of Total Weight, Shoot Weight, Root Weight, Primary Root Length, Root Weight per Length, and Shoot to Root Ratio of inbred lines

| Inbred Line | BVRB_3g058990 | BVRB_3g059000 | BVRB_3g059010 | BVRB_3g059020 | BVRB_3g059030 | BVRB_3g059040 | BVRB_3g059050 | BVRB_3g059060 | BVRB_3g059070 | BVRB_3g059080 | BVRB_3g059090 | BVRB_3g059100 |
|---|---|---|---|---|---|---|---|---|---|---|---|---|
| I | I/I | I/I | I/I | I/I | I/I | I/I | I/I | I/I | I/I | I/I | I/I | I/I |
| 55 | I/I | I/I | I/I | I/I | I/I | I/I | I/I | I/I | I/I | I/I | I/I | I/I |
| 57 | I/I | I/I | I/I | I/I | I/I | I/I | I/I | I/I | I/I | I/I | I/I | I/I |
| 59 | I/I | I/I | I/I | I/I | I/I | I/I | I/I | I/I | I/I | I/I | I/I | I/I |
| 66 | I/I | I/I | I/I | I/I* | I/I | I/I | I/I | I/I | I/I | I/I | I/I | I/I |
| 71 | I/I | I/I | I/I | I/I | I/I | I/I | I/I | I/I | I/I | I/I | I/I | I/I |
| G | G/G | G/G | G/G | G/G | G/G | G/G | G/G | G/G | G/G | G/G | G/G | G/G |
| 49 | G/G | G/G | G/G | G/G | G/G | G/G | G/G | G/G | G/G | G/G | G/G | G/G |
| 60 | G/G | G/G | G/G | G/G* | G/G | G/G | G/G | G/G | G/G | G/G | G/G | G/G |
| 78 | G/G | G/G | G/G | G/G | G/G | G/G | G/G | G/G | G/G | G/G | G/G | G/G |
| A | I/G | I/G | I/G | I/G | I/G | I/G | I/G | I/G | I/G | I/G | I/G | I/G |
| 52 | G/Z | G/Z | G/Z | G/Z | G/Z | G/Z | G/Z | G/G | G/G | G/G | G/G | G/G |
| 58 | Z/Z | Z/Z | Z/Z | Z/Z | Z/Z | Z/Z | Z/Z | G/G | G/G | G/G | G/G | G/G |
| H | I/D | I/D | I/D | I/D | I/D | I/D | I/D | I/D | I/D | I/D | I/D | I/D |
| 46 | B/X | B/X | I/X | I/X | I/X | I/X | I/X | I/X | I/X | I/X | I/X | I/X |
| 51 | G/B | G/B | G/B | G/B | G/B | G/B | G/B | G/B | G/B | G/B | G/B | G/B |
| 65 | I/Y | I/Y | I/Y | I/I* | I/Y | I/Y | I/Y | I/Y | I/Y | I/Y | I/Y | I/Y |
| F | F/F | F/F | F/F | F/F* | F/F | F/F | F/F | F/F | F/F | F/F | F/F | F/F |
| 50 | F/F | F/F | F/F | F/F* | F/F | F/F | F/F | F/F | F/F | F/F | F/F | F/F |
| 54 | F/F | F/F | F/F | F/F | F/F | F/F | F/F | F/F | F/F | F/F | F/F | F/F |
| 56 | F/F | F/F | F/F | F/F* | F/F | F/F | F/F | F/F | F/F | F/F | F/F | F/F |
| 77 | F/Y | F/Y | F/Y | F/Y | F/Y | F/Y | F/Y | F/Y | F/Y | F/Y | F/Y | F/Y |
| B | B/B | B/B | B/B | B/B | B/B | B/B | B/B | B/B | B/B | B/B | B/B | B/B |
| D | D/D | D/D | D/D | D/D | D/D | D/D | D/D | D/D | D/D | D/D | D/D | D/D |
| 63 | D/D | D/D | D/D | D/D* | D/D | D/D | D/D | D/D | D/D | D/D | D/D | D/D |
| I/G allele avg. TDW ± T22 | 0.25 ± 0.05 | 0.25 ± 0.05 | 0.25 ± 0.05 | 0.29 ± 0.06 | 0.25 ± 0.05 | 0.25 ± 0.05 | 0.25 ± 0.05 | 0.19 ± 0.06 | 0.19 ± 0.06 | 0.19 ± 0.06 | 0.19 ± 0.06 | 0.19 ± 0.06 |
| F allele avg. TDW ± T22 | -0.11 ± 0.12 | -0.11 ± 0.12 | -0.11 ± 0.12 | -0.11 ± 0.12 | -0.11 ± 0.12 | -0.11 ± 0.12 | -0.11 ± 0.12 | -0.11 ± 0.12 | -0.11 ± 0.12 | -0.11 ± 0.12 | -0.11 ± 0.12 | -0.11 ± 0.12 |
| *p*-value | 0.006 | 0.006 | 0.006 | 0.007 | 0.006 | 0.006 | 0.006 | 0.026 | 0.026 | 0.026 | 0.026 | 0.026 |
| F aa mutation | Yes | Yes | Yes | Yes | No | No | Yes | Yes | Yes | No | Yes | Yes |
| Exp. Seedling | Yes | Yes | Yes | Yes | Yes | Yes | No | Yes | No | Yes | No | Yes |
| Exp. Roots | Yes | Yes | No | Yes | Yes | Yes | No | Yes | Yes | Yes | No | No |

**Supplemental Figure 6: Summary of allele determination of loci within marker region.** Summary of the genome resequencing and Sanger sequencing (denoted by *) within inbred lines denoting loci in the marker region and allele specific genotyping I/G allele (blue), F allele (orange) or other (white) (dark is homozygous light is heterozygous). Genotypes identified through sequencing are identified as X, Y, or Z. Additional data includes marker data and total dried with ± standard error. average phenotypic trait by I/G or F allele types and associated *p*-value, F amino acid (aa) unique mutations within loci, and expression (Exp.) in either seedling or root tissue (FPKM≥0.5) (Dohm *et al.* 2014; Minoche *et al.* 2015) by loci.

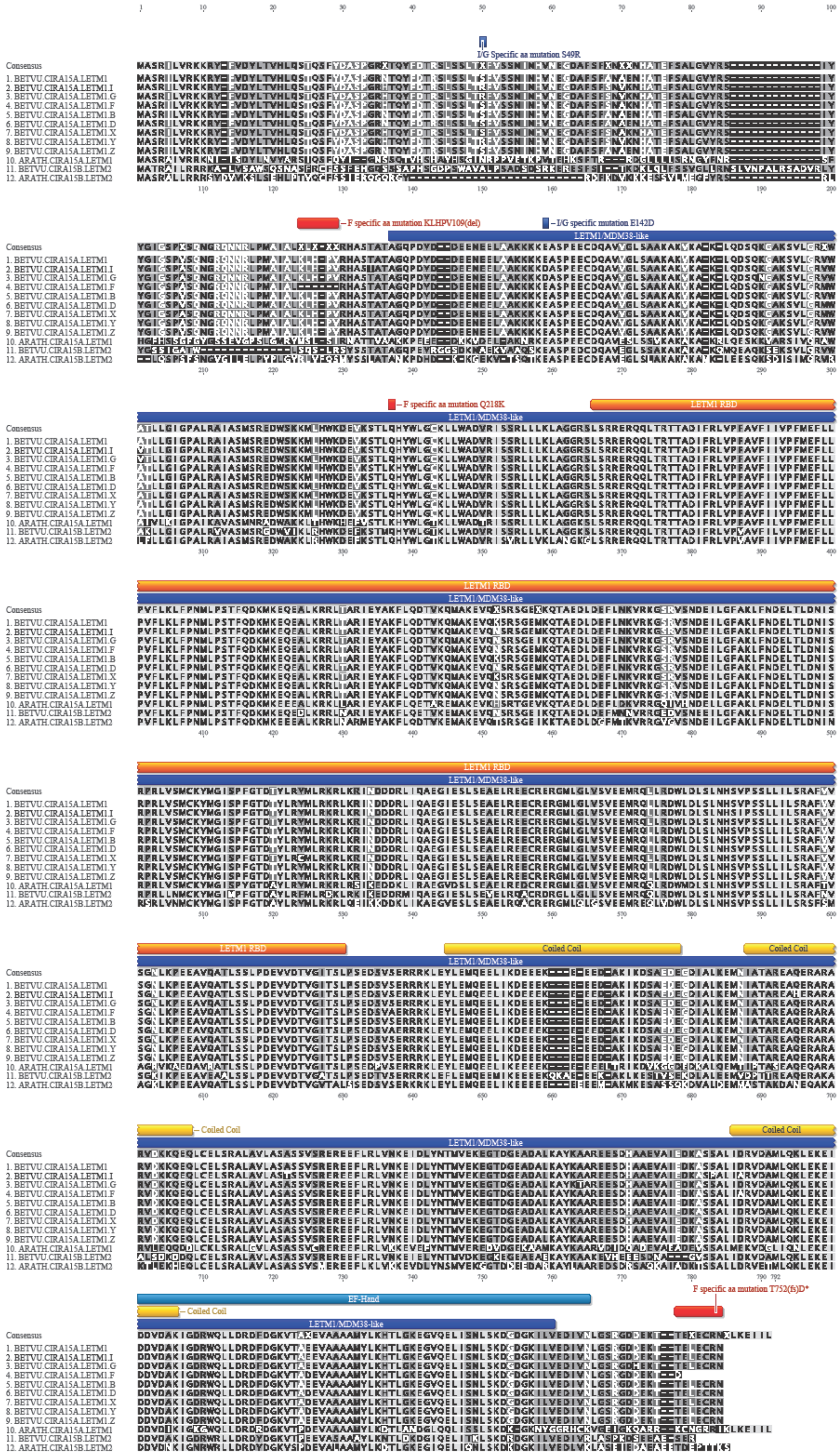

**Supplemental Figure 7: Mutational analysis of *BvCIRA15A/LETM1* alleles.** Amino acid alignment of *BvCIRA15A/LETM1* (BETVU.CIRA15A.LETM1) alleles identified in this study along with homolog *AtLETM1* (ARATH.CIRA15A.LETM1) and paralogs *BvCIRA15B* (BETVU.CIRA15B.LETM2), and *AtLETM2* (ARATH.CIRA15B.LETM2).

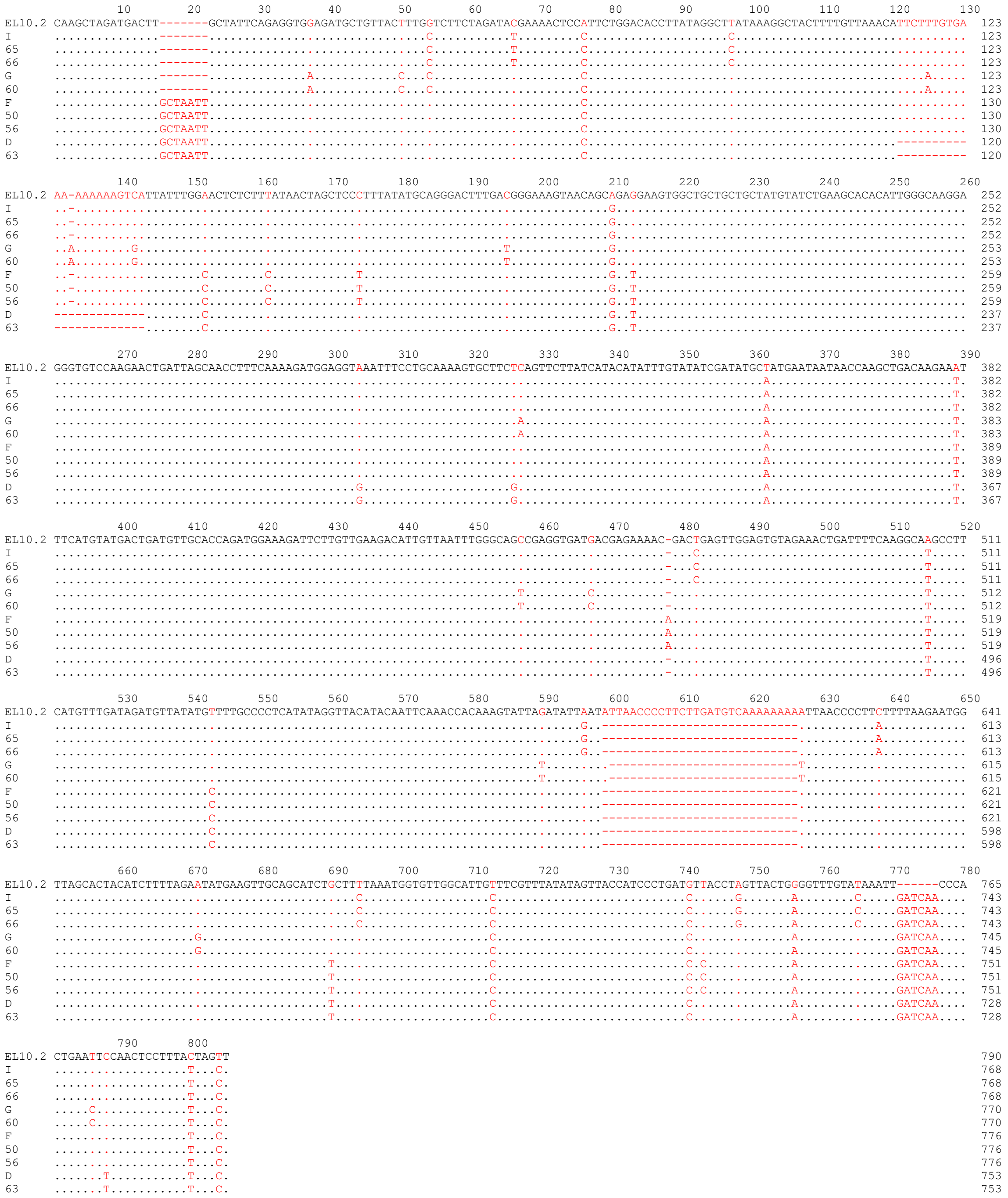

**Supplemental Figure 8) Summary of Sanger sequencing of *BVRB_3g059020* for allele determination.** Summary sequence data obtained on inbred lines with polymorphic alleles. Polymorphisms to EL10.2 sequence noted by red text.

```
                               10         20         30         40         50         60
CONSENSUS              XXXXXXXXXX XXXXXXXXXX XXXXXXXXXX MXXXRXXXXX XXXXXXXXXX XXXXXRRXXX
BVRB_3G058990 I ALLELE ---------- ---------- ---------- MNASRDRLSR PVDISSLLQN ---AERRVDL
BVRB_3G058990 F ALLELE ---------- ---------- ---------- MNASRDRLSR PVDISSLLQN ---AERRVDL
SOLYC10G080400.1.1     ---------- ---------- ---------- ---------- ---------- ----MGRGGI
AT3G57860              ---------- ---------- ---------- MPEARDRTER PVDYSTIFAN ----RRRHGI
OS02G0591500           ---------- ---------- ---------- ---------- ---------- ----------
OS04G0472700           MVSGFLRFGG PLFCPFFDEL FAGCSDASVC LCRCRSGEVF ESEQEAEEDF RGIGGEGTLA
MUSAC LOC103985512     ---------- ---------- ---------- M-------NQ GLGTMVDAGE -GFWIRTVAF
MUSAC LOC103989966     ---------- ---------- ---------- MPEAR----- -RATTGDFTG -GFWIRRAAY
MUSAC LOC103996371     ---------- ---------- ---------- MAETRQGVNR GRETTGSATN LGFWIRRAAY

                               70         80         90        100        110
CONSENSUS              XXXXXXXXXX XXSXXXXXXX XXXXXXXXXX XXXXXXXXXX XXXXXXXXXR ---XRRXXXX
BVRB_3G058990 I ALLELE VVDEPGLHWL GLSPQSVS-- ----TSQSGG TSGKSSRR-- --RVGLQKYR ---HRRYIS-
BVRB_3G058990 F ALLELE VVDEPGLHWL GLSPESVS-- ----TSESGG ASGKSSRR-- --RVGLQKYR ---HRRYIS-
SOLYC10G080400.1.1     EIFEDE---- --SPESSSRA PIQTTAEAGR MAGTSGGRGG IGRIGFGSPR ---NRRGRNL
AT3G57860              LLDEPD---- --SRLSLIES PVNPD----- -IGSIGGTGG LVRGNFTTWR PGNGRGGHTP
OS02G0591500           ---------- ---------- ---------- ---------- ---------- ----------
OS04G0472700           RIHTSRSKDQ REDQGYYSNM PEMRDSKRTA LGELSGGGGF FIRRVASPGA LAARGPGKPL
MUSAC LOC103985512     PDAVVR---- --S------- ---------- ---------- ---------R ---SRREQSL
MUSAC LOC103989966     PAAMPG---- --S------- ---------- ---------- ---------R ---RRRIQSL
MUSAC LOC103996371     LASRTG---- --S------- ---------- ---------- ---------R ---STRVQSV

                              120        130        140        150        160        170
CONSENSUS              XXXX---XBX ENXXPXXXXX XXXXXXXRXS XLPXWYPRTP LRDITXIVXA JERXRXXXXX
BVRB_3G058990 I ALLELE -RSL---LNQ ENQRPRSV-- --RKGQGRGS ILPSWYPRTP LRDITAIVRA IERKRAE---
BVRB_3G058990 F ALLELE -RSL---LNQ ENQRPRSV-- --RKGQGRGS ILPSWYPRTP LRDITAIVRA IERKRAE---
SOLYC10G080400.1.1     FRTPARVIGR QNISPTGQ-- --GRNRGRHS VLPAWYPRTP LRDITSIVRA IERTRAR---
AT3G57860              FRLP---QGR ENMPIVTA-- --RRGRG-GG LLPSWYPRTP LRDITHIVRA IERRRGA---
OS02G0591500           ---------- ---QYAYNFT YHLQAVERKS RLGN----AA VRQ--QIQLS EDSSRSVDPA
OS04G0472700           ARRFIRPSNN KENVPPV-WA VKATATKRRS PLPDWYPRTP LRDITAIAKA IQRSRLRI-A
MUSAC LOC103985512     VFTT---DDK ENIPPSLVVT ARRRVGRRKS PLPSWYPRTP LRDVTVVVNA LERRRMRAT-
MUSAC LOC103989966     VAAI---DDK ENIHPSRAAT ARRRPRNRKS PLPSWYPRTP LRDITIIVNA LER-RMRER-
MUSAC LOC103996371     VFAT---DDK ENVTPSRA-- ARRRGRTRKS PLPEWYPRTP LRDITVIVNA LERRRMRVRA

                              180        190        200        210        220        230
CONSENSUS              XXXXXXXXXX XXXXXXXXXX XXXXXXXXXX XXXXXXXXXX XXXXXX-XXX XXXXXSXXXX
BVRB_3G058990 I ALLELE -----LRDND ---------- -QMIESPQQ- ETNSLDTTS- ---------E LNNEMSISTP
BVRB_3G058990 F ALLELE -----LRDND ---------- -QMIESPQQ- ETNSLDTT-- ---------D LNNEMSISTP
SOLYC10G080400.1.1     -----LRESE GEQ------- -LESVVPQDH TDLGPSESTS -------GAQ LEHTNSLITP
AT3G57860              -------GTG GDD------- GRVIEIPTHR QVGVLESPV- -------PLS GEHKCSMVTP
OS02G0591500           TPVQK----- EEGVPQSTPT PPTQKALDAA APCPGSTQAV ASTSTAYLAE GKPKASSSSP
OS04G0472700           AAQQR----- SQTPEQNTPH CTEVRDSLDV EPGINSTQIV ATPASS-LAK DSLKI-FSSP
MUSAC LOC103985512     AARVKRRNRN ---------- -REMKEPDQT SLGEALSSDV ---------- -SDAWYVSPI
MUSAC LOC103989966     AARARQRNAN PEASPVTDPA VDEMNASEHT PVGEALPSDV SSDASPSPAA LSTLSSLTPT
MUSAC LOC103996371     AATARQRTSD PE-------- ------PAAV EKGLLDSSSI -------PAA AGS-SSVSAT

                              240        250        260        270        280        290
CONSENSUS              XXXXXXXXXX XXXXXXXXXX XXXXXXXXXX XXX--XXXXX ZXXLXXXIXX XEXXVXXXXX
BVRB_3G058990 I ALLELE PQACKFK--- ---------- --DDNASKKG KDI--EFFTP QKKLLNSIEK VRHIWLEDQR
BVRB_3G058990 F ALLELE PQACKFK--- ---------- --DDNASKKG KDI--EFFTP QKKLLNSIEK VRHIWLEDQR
SOLYC10G080400.1.1     RPKTR-SRYH TRSVGKVPKI LLDITNQSTS EDA--ECLTP QRKLLNSIDT VEKHVMEELH
AT3G57860              GPSVGFKRSC PPSTAKVQKM LLDITKEIAE EEA--GFITP EKKLLNSIDK VEKIVMAEIQ
OS02G0591500           SD-CSFQTPS RP----NDPA LADLME---- ---------- -KELSSSIEQ IEKMVRKNLK
OS04G0472700           SE-TSLVTPS KP----MDPV LLDDME---- ---------- K-KLSSSIEQ IEKMVRRNLK
MUSAC LOC103985512     GHSLQSPSSG LVSVS----- -----TDPST EDM--RPTEF EERLQSTIAE MERLVLRNLK
MUSAC LOC103989966     QQPLRTPSSS ETPLSTTERV L----ADPST EDP--KPTEF EKKLQSTISE MERLVLDNLK
MUSAC LOC103996371     EQPPQICSSS N--------- -----ASSPR EDPPDQPTEY EKNLEIYIGE MERLVTENLK

                              300        310
CONSENSUS              XXXXXXAXXX XXXXXXXRTL XSMR
BVRB_3G058990 I ALLELE KLERTPAAKK AERKKMVRVL MSMR
BVRB_3G058990 F ALLELE KLERTPAAKK AERKKMVRVL MSMR
SOLYC10G080400.1.1     KLKRTPSARK QERDKRVKTL MSMR
AT3G57860              KLKSTPQAKR EEREKRVRTL MTMR
OS02G0591500           RAPK--AAQP SKVTIQKRTL LSMR
OS04G0472700           RTPKAAAAQP SKRAIQRRTL MSMR
MUSAC LOC103985512     RSPEPHA--- -KMKKPTRTL LSMR
MUSAC LOC103989966     RTPKPPA--- ---RKAMRTL MSMR
MUSAC LOC103996371     RSPMPPA--- ---KRAKRTL ISMR
```

**Supplemental Figure 9: Alignment of BVRB_3g058990 mutations found in inbred line F**
Amino acid alignment of *BVRB_3g058990* alleles from study and homologs from tomato (SOLYC), *Arabidopsis* (AT), rice (OS), and banana (MUSAC). Alignment performed by MUSCLE (Edgar 2004), consensus set to 75% shared identity. Mutated sequence and corresponding aligned sequence identified by blue (non-F specific) and red (F specific) text.

```
                                          10         20         30         40         50         60
CONSENSUS                                GEKGXG-XXG XXXXYKGXX FHRXIXXFMXQ GGDXXXGBGX GGXSIY---G XXFXDENFXX
BVRB_3G059000 I ALLELE 130 GEKGYG---- ----YKGCH FHRIIKDFMIQ GGDFTEGNGT GGISIY---G SKFEDENFTL 178
BVRB_3G059000 F ALLELE 130 GEKGYG---- ----YKGCH FHRIIKDFMIQ GGDFTEGDGT GGISIY---G SKFEDENFTL 178
AT2G38730               70 GELR---KAG KPLGYKECQ FHRVIKDFMVQ SGDFLKNDGS GCMSIY---G HKFEDENFTA 123
SOLYC01G096520          61 GEFR---KAG VPQGFKNCQ FHRVIKDFMIQ GGDFLKGDGS GCVSIY---G SKFEDENFIA 114
MUSAC CAG1860793        80 GEYR---KAN LPIGYKGCQ FHRIIKDFMIQ AGDFLKGDGS GCVSIY---G SKFEDENFIA 133
OS03G0811600            75 GEHR---KSG LPQGYKGCQ FHRVIKDFMIQ GGDYMKGDGT GCTSIY---G TKFDDENFIA 128
AT3G62030              186 GEKKYG---- ----YKGSS FHRIIKDFMIQ GGDFTEGNGT GGISIY---G AKFEDENFTL 234
OS06G0216800            64 GEKGVGAVTG KHLHYKGSC FHRVIKGFMVQ GGDITAGDGT GGESIY---G LKFEDENFVL 120
OS02G0761100            71 GEKGVSAATG VPLHYKGSC IHRIVKGFMVQ GGDITAGDGT GGESIY---G LNFEDENFVL 127
MUSAC CAG1832565        54 REKGVGPHTG VPLHFKGSC FHRVIKGFMIQ GGDISAGDGT GGESIY---G LKFEDENFIL 110
MUSAC CAG1837590        60 GEKGVGPNTG VPLHLKGSC FHRVIKGFMIQ GGDISAGDGT GGESIY---G LKFEDENFAL 116
SOLYC01G108340          42 GEKGIGPNTG VPLHYKGNC FHRVIKSFMVQ GGDISAGDGT GGESIY---G LKFEDENFEL  98
BVRB_6G134480           43 GEKGISPISG RSLHYKGMR FHRIVKGALIQ GGDISGEGES GGESIY---G PTFDDENFEL  99
AT1G01030               42 GEKGLGPNTG VPLHYKGNR FHRVIKGFMIQ GGDISANDGT GGESIY---G LKFDDENFEL  98
AT2G15790               42 GEKGLGPNTG VPLHYKGNR FHRVIKGFMIQ GGDISANDGT GGESIY---G LKFDDENFEL  98
BVRB_6G150350          131 GEKGFG---- ----YKGSS FHRVIKDFMIQ GGDFDKGNGT GGKSIY---G RTFKDENFNL 179
AT5G13120              131 GEKGFG---- ----YKGST FHRVIRDFMIQ GGDFEKGNGT GGKSVY---G RTFKDENFKL 179
MUSAC CAG1846609       120 GEKGFG---- ----YKGSA FHRVIKDFMIQ GGDFDQGNGT GGKSIY---G RTFKDENFKL 168
MUSAC CAG1862883       123 GEKGFG---- ----YKGSA FHRVIKDFMIQ GGDFDKGNGT GGKSIY---G RTFKDENFKL 171
OS05G0103200           124 GEKGFG---- ----YKGSS FHRVIKDFMIQ GGDFDKGNGT GGKSIY---G RTFKDENFKL 172
SOLYC01G009990         122 GEKGFG---- ----FKDSA FHRVIKDFMIQ GGDFDKGNGT GGKSIY---G RTFKDENFKL 170
AT3G56070               42 GENGIG-KAG KALHYKGSA FHRIIPGFMCQ GGDFTRGNGT GGESIY---G SKFEDENFKL  97
BVRB_4G095580           44 GEKGLG-KLG KPLHYKGSA FHRIIPNFMCQ GGDFTRGNGT GGESIY---G SKFDDENFKL  99
OS09G0571400            44 GEKGLG-ASG KPLHYKGSA FHRIIPNFMCQ GGDFTRGNGT GGESIY---G DRFADENFKL  99
OS02G0121300            42 GEKGVG-KSG KPLHYKGST FHRVIPEFMCQ GGDFTRGNGT GGESIY---G EKFADEVFKF  97
MUSAC CAG1849936        42 GEKGMG-RCG KPLHYKGSS FHRVIPGFMCQ GGDFTRGNGT GGESIY---G EKFADENFVK  97
MUSAC CAG1859271        43 GEKGVG-RSG KPLHYKGST FHRVIPGFMCQ GGDFTCGNGT GGESIY---G EKFADENFVK  98
SOLYC01G111170          42 GEKGVG-KMG KPLHYKGST FHRVIPGFMCQ GGDFTAGNGT GGESIY---G AKFNDENFVK  97
AT4G38740               42 GEKGVG-GTG KPLHFKGSK FHRVIPNFMCQ GGDFTAGNGT GGESIY---G SKFEDENFER  97
AT2G21130               43 GEKGVG-RSG KPLHFKGSS FHRVIPNFMCQ GGDFTKGNGT GGESIY---G AKFEDENFER  98
AT2G16600               43 GERGIG-KQG KPLHYKGSS FHRVIPKFMCQ GGDFTAGNGT GGESIY---G SKFKDENFIK  98
AT4G34870               42 GEKGMG-KLG KPLHFKGSI FHRVIPGFMCQ GGDFTAKNGT GGESIY---G AKFKDENFIK  97
OS10G0154700            49 GERAGR-SGK SRLHYKGSA FHRVVPGFMCQ GGDITAGNGT GGESALDGAA RHFPDEGFAV 107
AT3G22920               42 GERGIG-KCG KPIHYKGST FDHIVPDLMWC GGDIIFEN-- --EPIH---S EELDDEYFIL  93
MUSAC CAG1859024        46 GERGVE-RSG KPLHYKGTR FLRVTPGFMCQ GRDVAHNDGT GMESIY---G GKFADENFTT 101
OS09G0537600            76 GEEGIG-HKG KSLHYKGSR FHRIIPGFMIQ GGDIVRGDGK GSESIY---G GTFPDENFIV 131
MUSAC CAG1858381        76 GEKSNG-TGG RNLHYKGTK FHRIVSGFVIQ GGDITNGDGR GSESIY---G GTFPDENFVI 132
AT4G34960               85 GEKGKT-SSG KPLHYKGTP FHRIISGFVIQ GGDIIHGDGK SSDSIY---G GTFPDENFKI 140
OS06G0708400            78 GEKGLG-KSG KPLSYKGTP FHRIIPGFMIQ GGDTVSGNGT GCDSIY---G GMFPDENFKI 133
MUSAC CAG1855656        71 GEKGRD-KYG VRLYYKGST FHRIIPHFMIQ GGDIVFGNGR GIDNIY---D EPFADENFKL 126
AT2G29960               70 GEKGVG-KSG KPLHYKGSK FHRIIPSFMIQ GGDFTHGNGM GGESIY---G QKFADENFKL 125
AT5G58710               73 GEKGIG-KNG KALHYKGSS FHRIIPSFMLQ GGDFTHGNGM GGESIY---G EKFADENFKL 128
SOLYC06G051650          94 GEKGTG-KAG KPLHYKGSS FHRIIPSFMIQ GGDFTRGDGR GGESIY---G ESFPDENFDL 149
AT3G55920               97 GEKGVG-NMG KPLYFKGSS FHRIIPGFMIQ GGDFTRGDGR GGESIY---G DKFADENFKL 152
BVRB_7G172160           74 GEKGVS-KSG KPLHYKGST FHRIIPSFMIQ GGDFTRGDGT GGESIY---G EKFADENFKL 129
MUSAC CAG1852830        76 GEKGVG-KSG KALHFKGST FHRIIPSFMIQ GGDFTHGDGR GGESIY---G MTFADENFKL 131
MUSAC CAG1856974        85 GEKGIG-KSG KTLHYKGSA FHRIIPSFMIQ GGDFTLGDGR GGESIY---G SKFADENFKL 140
OS06G0708500            94 GEKGTG-KSG KALHFKGSA FHRIIPSFMIQ GGDFTLGDGR GGESIY---G TKFADENFKI 149
```

**Supplemental Figure 10: Alignment of BVRB_3g05900 mutation found in inbred line F**
Amino acid alignment of *BVRB_3g05900* alleles from this study and homologs from tomato (SOLYC), *Arabidopsis* (AT), rice (OS), and banana (MUSAC). Alignment performed by MUSCLE(Edgar 2004), consensus at 75% shared identity. Bold text represents amino acid number in original sequence, mutated sequence and corresponding aligned sequence identified by red text.

```
                                   180        190        200        210        220        230        240
Consensus                          WEEYXXIXXX -XXXXFKIX- HXXXPLSYYT GJLGMXGXTA YXGFXEXXXP KKGXXVXVSA
BVRB_3g059060 I Allele 107         WEEYSLISSP SPDMFFKIE- HTDVPLSYYT GILGMPGITA YGGFYELASP KKGDLVYVSA   166
BVRB_3g059060 F Allele 107         WEEYSLISSP SPGIFFKIE- HTDVPLSYYT GILGMPGITA YGGFYELASP KKGDRVYVSA   166
BVRB_4g074550          160         WEEYSLIEDT --KKLRKIQ- QDDIPLSYHI GLLGMPGFTA YAGFYEVCSP KEGDRVFVSA   217
BVRB_3g059080          108         WEEYTLTTLT --DLFFKIE- HTDVPLSYYT GLLGMAGMTA YVGFYELGSP KKGDRVYVSA   167
AT3G59845              110         WEEFSTINP- --AAIFKIDV NINVPLSYYT GILGMIGLTA YAGFFEICSP KKGDTVFVSA   167
AT5G16960              107         WEEYSVITPI -PNLHFKIH- HTNFPLSYYT GLLGMPGMTA YVGFYEICTP KKGDTVFVSA   165
AT5G37940              114         WEEYSVITPT -PSSHFKIH- HTDVPLSFYT GLLGIPGLTA YVGFYEICSP KKGETVFVSA   172
AT5G38000              114         WEEYSVITPT -PSSHFKIH- HTDVPLSFYT GLLGIPGLTA YIGFYEICSP KKGETVFVSA   172
AT1G26320              112         WEEYSVITLT -TYSHFKIE- HTDVPLSYYT GLLGMPGMTA YAGFYEVCSP KKGETVFVSA   170
AT3G03080              112         WGEYSLITPD --FSHYKIQ- HTDVPLSYYT GLLGMPGMTA YAGFYEICSP KKGETVFVSA   169
AT5G16980               74         -----LT--- --HEHFKIQ- HTDVPLSYYT GLLGMPGMTA YVGFYEICSP KKGETVYVSA   124
AT5G16970              106         WEEYSVITPM -THAHFKIQ- HTDVPLSYYT GLLGMPGMTA YAGFYEVCSP KEGETVYVSA   164
AT5G17000              106         WEEYSVITPM -THMHFKIQ- HTDIPLSYYT GLLGMPGMTA YAGFYEVCSP KEGETVYVSA   164
AT5G16990              104         WEEYSVITPM -AHMHFKIQ- HTDVPLSYYT GLLGMPGMTA YAGFYEVCSP KKGETVYVSA   162
Os04g0497000           106         WEEYSLIKDP -SRALFAIR- HPDLPLSYYT GLLGMAGFTA YVGFHEICAP REGERVYVSA   164
Os11g0255500           120         WEEYSLITQP --ETLHKIN- HPDLPLSYYT GVLGVTGLTA YAAFFEVGKP KKGETVFVSA   177
Os12g0225900           106         WEEYTLVNNP KP-YLHKIN- YPEFPLSYYT GVLGIAGLTA YGGFFEVSKP KKGDYVFVSA   164
Os12g0226400           111         WEEYTLINNP --ESLFKIN- YPEFPLSNYT GVLGMHGLTA YVGFFEMSKP KKGEYVFVSS   168
Os12g0227400           108         WEEYTVINNP --ETLFKIN- HPELPLSYYT GILGMPGLTA YGGFFEVAKP KKGEYVFISS   165
Os12g0226900           108         WEEYTVINNP --ESLFRIN- HPKLPLSYYT GILGMPGLTA YAGFFEVSKP KKGEYVFISA   165
Os12g0226700           108         WEEYTVINNP --ESLFKIN- HPELPLSYYT GILGMPGLTA YAGFFDVPKP KKGEYVFISG   165

                                  300        310        320        330        340        350        360
Consensus                          PXGIDIYFEN VGGXMLDAVL XNMXXXGRIA XCGMISQYNL XXXEGVXNLX XIXXKRJRXZ
BVRB_3g059060 I Allele  225        PEGIDIYFEN VGGKMLEAVL SNMRVKGRIA VCGMISQYNL EQPEGVHNLF HITTKRIRME   284
BVRB_3g059060 F Allele  225        PEGIDIYFEN VGGKMLEAVL SNMRVKGRIA VCGMISQYNL EQPEGVHNLF QITTKRIRME   284
BVRB_4g074550           276        PEGIDIYFEN VGGAMLDAVL LNMRAHGRIA VCGMVSQHSN HDKAGIHNLF TLIRNRVTMK   335
BVRB_3g059080           224        PEGIDIYFEN VGGKMLEAVL SNMRNNGRIP VCGMISQYNL VEPEGVHNLF NITTKRIRME   283
AT3G59845               226        PKGIDIYFEN VGGKMLDAVL LNMKTYGRIA VCGMISQYHL ETRDRLQNLP DIIFKKIRMQ   285
AT5G16960               224        PEGIDIYFEN VGGKMLDAVI LNMRPHGRIA ACGMISQYNL KNPEGIYGLS LITYKRIRIE   283
AT5G37940               231        PEGIDIYFEN VGGKMLDAVL QNMRTHGRIA ACGMISQYNL KEPEGLHNTA TIVHKRIRVQ   290
AT5G38000               231        PEGIDIYFEN VGGKMLEAVL ENMRTHGRIA ACGMISQYNL KKPEVLHNTA TIVHKRIRVQ   290
AT1G26320               229        PEGIDIYFEN VGGKMLDAVL INMKLHGRVA VCGMISQYNL VDPEGVHNLP TILYKRIQLQ   288
AT3G03080               228        PEGIDIYFEN VGGKMLDAVL LNMKLHGRIA VCGMISQYNL EDQEGVHNLA NVIYKRIRIK   287
AT5G16980               183        PNGIDIYFEN VGGKMLDAVL MNMNMHGRIA VCGMISQYNL ENQEGVHNLS NIIYKRIRIQ   242
AT5G16970               223        PNGIDIYFEN VGGKMLDAVL VNMNMHGRIA VCGMISQYNL ENQEGVHNLS NIIYKRIRIQ   282
AT5G17000               223        PTGIDIYFEN VGGKMLDAVL LNMNPHGRIA VCGMISQYNL ENQEGVHNLS NIIYKRIRIQ   282
AT5G16990               221        PKGIDMYFEN VGGKMLDAVL LNMNPHGRIA VCGMISQYNL ENQEGVHNLS NIIYKRIRIQ   280
Os04g0497000            223        PEGIDIYFEN VGGAMLDAVL LNMRVRGRVA ACGMISQYNL EHPDPVHNLT AIVTKRLRIE   282
Os11g0255500            237        PDGIDIYFDN VGGATLDAAL VNMRRGGRVV VCGMISQYNL QEPEGVHNVI QILSKTIRVE   296
Os12g0225900            223        PDGIDIYFDN VGGAMLDAVL PNMRIGGKIT ICGMISQYNL ERPDGVRNLF YLFAKSLRME   282
Os12g0226400            227        PEGIDIDFEN VGGAMLDAVL PNMRLGGRIT MCGMISQYHL ERPEGVRNLM YIITKRLRME   286
Os12g0227400            224        PDGIDIYFEN VGGAMLDAVL PNMRVAGRIA ACGMISQYNL EQPEGVYNMI CIVTKRLRMQ   283
Os12g0226900            224        PEGIDIYFEN VGGETLDAVL PNMRLGGRIA ACGMISQYNL ERPEGVKNLF YIVTKRLRME   283
Os12g0226700            224        PEGIDIYYEN VGGKMLDAVL PNMSLGGRIV ACGMISQYNL EQPEGVRNLY YIVTKRLRME   283
```

**Supplemental Figure 11: Alignment of *BVRB_3g059060* mutation found in inbred line F**
Amino acid alignment of *BVRB_3g059060* alleles from study and homologs from *Beta vulgaris* (BVRB), *Arabidopsis* (AT) and rice (OS). Alignment performed by MUSCLE (Edgar 2004), consensus set to 75% shared identity. Mutated sequence and corresponding aligned sequence identified by blue (non-F specific) and red (F specific) text.

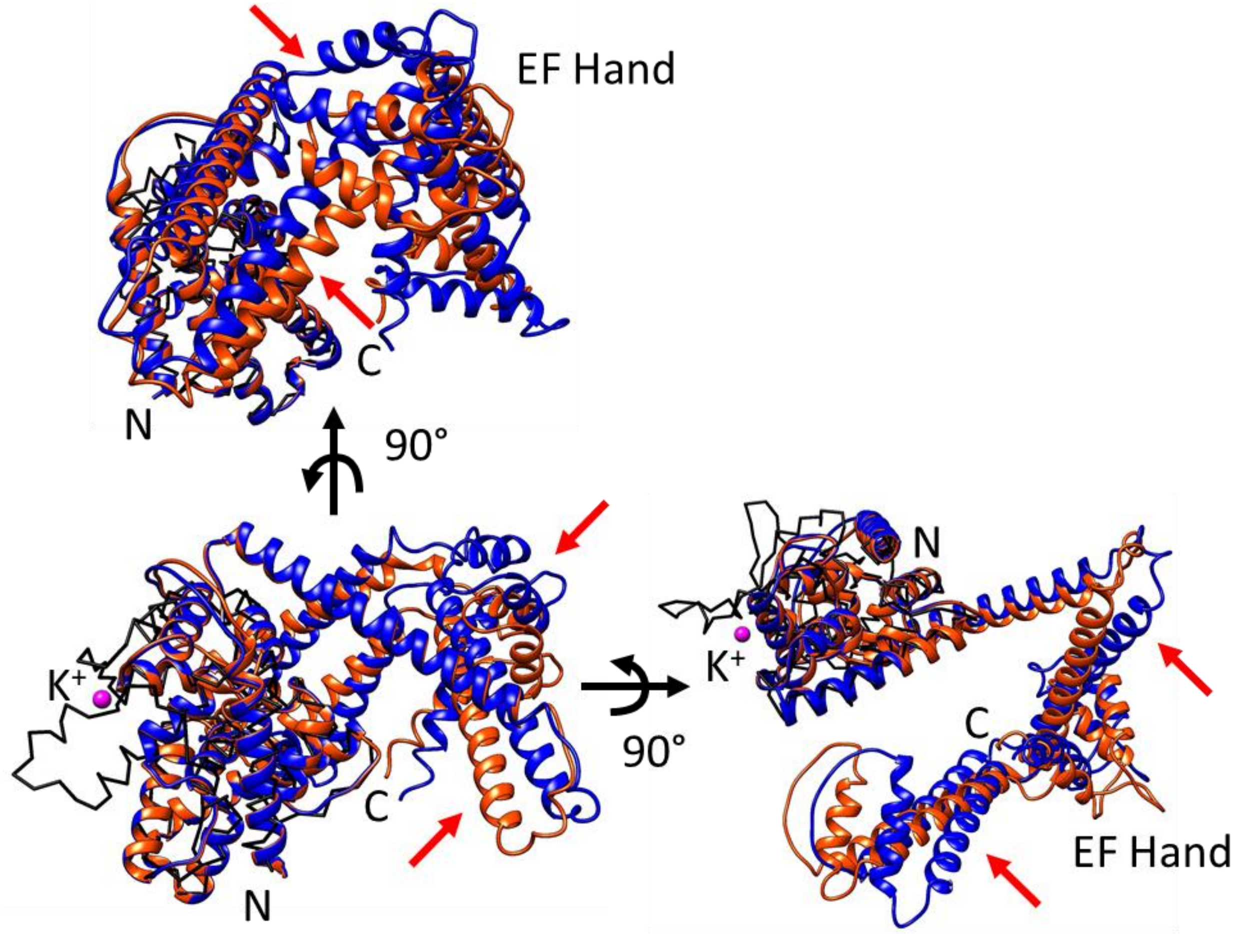


**Supplemental Figure 12: Prediction of structural effects of polymorphisms in BvCIRA15A/LETM1A.** Structures of the C-terminal parts of BvCIRA15A alleles were predicted using Alphafold2. Allele I/G is shown as blue ribbon (residues 308-758) and allele F is shown as red ribbon (residues 303-747). For comparison, the structurally determined ribosome-binding domain of the *Saccharomyces cerevisiae* MDM38 (pdb: 3skq) is shown as black chain trace, and the bound $K^+$ in MDM38 is shown in magenta. Red arrows point to regions of high allelic differences.

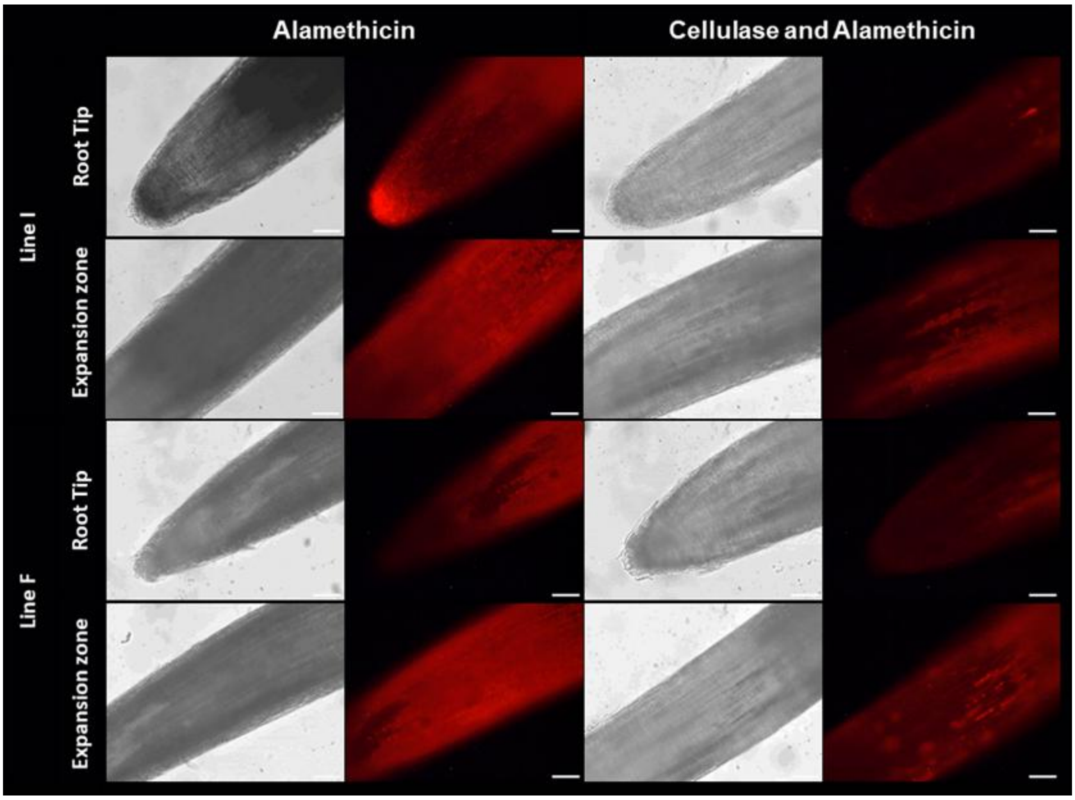


**Supplemental Figure 13: CIRA assay on sugar beet inbred lines I and F.** Sugar beet seedling roots from line I and F were assayed for CIRA competence. Three to five-day old seedlings from each line were subjected to control or cellulase (2-hour, 1% *T. viride* cellulase) followed by 10 min treatment with 20 µg/mL Alamethicin treatment. All were then stained with 1 µM of fluorescent DNA probe propidium iodide for 1 minute. Both bright field and fluorescent images were captured. Positive staining indicates penetration of alamethicin into cell membranes and the formation of open channels allowing for the intercalation of propidium iodide intracellularly. The two lines are different from each other in susceptibility to alamethicin and line I shows a clear CIRA response in the root tip with reduced fluorescence after cellulase pretreatment, whereas line F has a milder phenotype. Representative images of 3 independent CIRA assays with pools of at least 3 seedlings. Bars represent 100 µm.

```
                   10        20        30        40        50        60        70        80        90       100
Consensus TGTCGTCTCGTCTGAACCACGTCAATGTTCACACCGGATGGGGGAAAGTGTATTAGTTTAGCATCTTTTGTATGCTTTAGGGCCATCCATGATAAAATTT   100
Barke     ....................................................................................................   100
Bonus     ....................................................................................................   100
Morex     ....................................................................................................   100
FT880     ....................................................................................................   100

                  110       120       130       140       150       160       170       180       190       200
Consensus GTTCCATGAGATCTATTTGCCCAGTTTTAGACTTTTAGTTCATAAATACCCAGTAAGTAGGTATTGGGATTTATGATTGTCTGGACCAGATTATTTTGTT   200
Barke     ....................................................................................................   200
Bonus     ....................................................................................................   200
Morex     ....................................................................................................   200
FT880     ....................................................................................................   200
                  210       220       230       240       250       260       270       280       290       300
Consensus GTTTCCAACATAATGCTGCGAACATTGAGCTGTTATGCTGCTAGTATCAAACTGTTACCCTGTTATTTATTTCCAACATAATGCTACTTTACCGCTAACA   300
Barke     ....................................................................................................   300
Bonus     ....................................................................................................   300
Morex     ....................................................................................................   300
FT880     ....................................................................................................   300
                  310       320       330       340       350       360       370       380       390       400
Consensus CTAAACTGTTGCCTGTTTATTTTATCTGTAGCATACTGCTGGTAATAGTGCCTGTAACCCAGTTATTTTCTTTTCAGCGTAGTGCTGCTTGTACCGAATT   400
Barke     ....................................................................................................   400
Bonus     ....................................................................................................   400
Morex     ....................................................................................................   400
FT880     ....................................................................................................   400
                  410       420       430       440       450       460       470       480       490       500
Consensus GTAACCAGCTATTTTATTTTCAGCGTAATGCTGCTTATGCCGAACTGTTGTTAACACTTTTGCATACAAAAATATTCAGGAGGAAGAGAAGAAGAAAGAG   500
Barke     ....................................................................................................   500
Bonus     ....................................................................................................   500
Morex     ....................................................................................................   500
FT880     ..............................................................................................----..   496

                  510       520       530       540       550       560       570       580       590       600
Consensus AAAGAAGAAAAGGCAAAACAAAAGAAGGAAGAGAAGGCAAAGCTCAAAGAACCAGAGGCTGATGAACAGGATTTAGCTTTAAAGGAAATGACTGAGCCTA   600
Barke     ....................................................................................................   600
Bonus     ....................................................................................................   600
Morex     ....................................................................................................   600
FT880     ....................................................................................................   596

                  610       620       630       640       650       660       670       680       690       700
Consensus CTGCTAGGGAAGAAGAAGAACTCAGAGAAGGAAAGCAGCATGATAAGGAACAACTCTGTAATATCAGTCGAGCATTGGCTGTGCTAGCATCTGCATCGGT   700
Barke     ....................................................................................................   700
Bonus     ....................................................................................................   700
Morex     ....................................................................................................   700
FT880     ....................................................................................................   696

                  710       720       730       740       750       760       770       780       790       800
Consensus ATGATCCTAGTC-TATCTGTAATTTATATTAGCACATTGACTACTGACAGTAATAGTCGATGCTTACTTATATTGCAGTCTGTTAGCAAGGAGCGTCAAG   799
Barke     ............T.......................................................................................   800
Bonus     ............-.......................................................................................   799
Morex     ............-.......................................................................................   799
FT880     ............-.......................................................................................   795

                  810       820       830       840       850       860       870       880       890       900
Consensus AGTTCCTGAGCCTTGTCAACCAGGAGGTACAGTGTGGACTCATGAAATGGAATTAGAGGAACAATTATCCGTACTTCGTTCTTCTGTTATGTTGCATATG   899
Barke     ....................................................................................................   900
Bonus     ....................................................................................................   899
Morex     ....................................................................................................   899
FT880     ....................................................................................................   895

                  910       920       930       940       950       960       970       980       990      1000
Consensus CTTGTACTGTGGAGGTCTAATAGGACTTTMTTATTGTGGAATGCAGATAGGACTGTATAACTCCATGATTGAAAAAGAAGGCACAGATGGTGAAGAAGCT   999
Barke     ....................................................................................................  1000
Bonus     ....................................................................................................   999
Morex     ....................................................................................................   999
FT880     ....................................................................................................   995

                 1010      1020      1030
Consensus AAGAGGGCATACATGGCTGCTAGAGAGGAGGC                                                                      1031
Barke     ................................                                                                      1032
Bonus     ................................                                                                      1031
Morex     ................................                                                                      1031
FT880     ................................                                                                      1027
```

**Supplemental Figure 14: Summary of Sanger sequencing of *HvLETM1-like2* for allele determination.** Summary sequence data obtained on Barley lines. Polymorphisms to MorexV3 sequence noted by red text, polymorphisms in intron encoding sequence denoted in blue text.

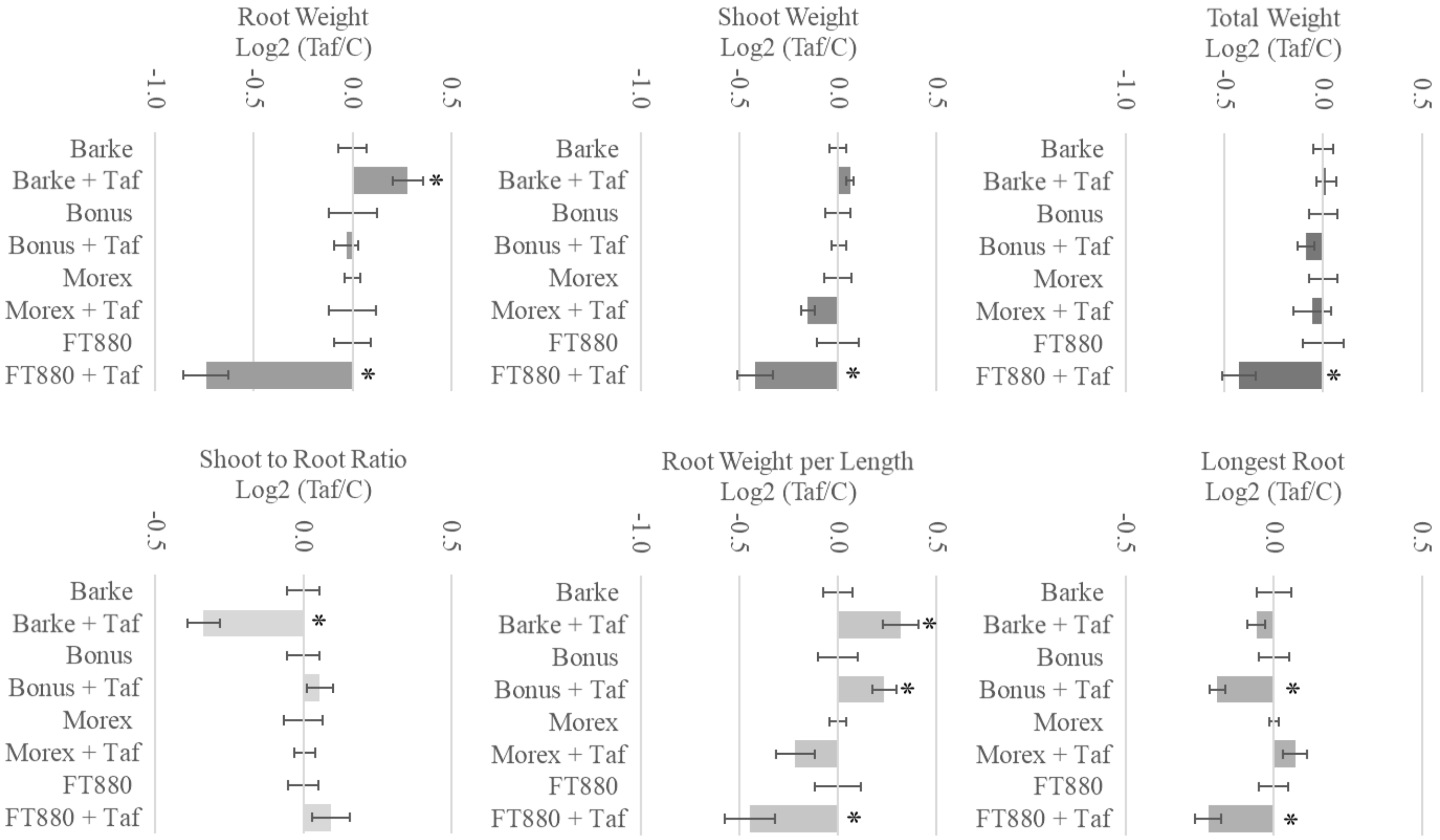


**Supplemental Figure 15: Phenotypic results of Barley lines.** Associated Barley lines of Barke, Bonus, Morex and HvLETM1-like2 mutated line FT880 with control and Taf treated plants. The log2 transformed and normalized to control phenotypes of Total Weight, Shoot Weight, Root Weight, Primary Root Length, Root Weight per Length, and Shoot to Root Ratio. Each line is the average of 4 biological replicates of at least 3 plants. Bars represent standard error. Asterisk represent significant difference between control and Taf treated total weight using an FDR *q* value of 0.05. (see Fig. S15).

| Line | Treatment | N | Total Weight Log2 Norm to Control | Shoot Weight Log2 Norm to Control | Root Weight Log2 Norm to Control | Longest Root Length Log2 Norm to Control | Root Weight per Length Log2 Norm to Control | Shoot to Root Ratio Log2 Norm to Control |
|---|---|---|---|---|---|---|---|---|
| Barke | C | 18 | 0.00 ± 0.05 | 0.00 ± 0.04 | 0.00 ± 0.07 | 0.00 ± 0.06 | 0.00 ± 0.07 | 0.00 ± 0.05 |
| Barke | Taf | 17 | 0.02 ± 0.05 | 0.06 ± 0.02 | **0.28 ± 0.08** | -0.06 ± 0.03 | **0.32 ± 0.09** | **-0.34 ± 0.05** |
| Bonus | C | 17 | 0.00 ± 0.07 | 0.00 ± 0.06 | 0.00 ± 0.12 | 0.00 ± 0.05 | 0.00 ± 0.1 | 0.00 ± 0.06 |
| Bonus | Taf | 18 | -0.09 ± 0.04 | 0.00 ± 0.04 | -0.03 ± 0.06 | **-0.19 ± 0.03** | **0.24 ± 0.06** | 0.06 ± 0.04 |
| Morex | C | 12 | 0.00 ± 0.07 | 0.00 ± 0.07 | 0.00 ± 0.04 | 0.00 ± 0.02 | 0.00 ± 0.04 | 0.00 ± 0.07 |
| Morex | Taf | 13 | -0.05 ± 0.1 | -0.15 ± 0.04 | 0.00 ± 0.12 | 0.07 ± 0.04 | -0.22 ± 0.1 | 0.00 ± 0.04 |
| FT880 | C | 13 | 0.00 ± 0.1 | 0.00 ± 0.11 | 0.00 ± 0.09 | 0.00 ± 0.05 | 0.00 ± 0.12 | 0.00 ± 0.05 |
| FT880 | Taf | 12 | **-0.43 ± 0.09** | **-0.42 ± 0.09** | **-0.74 ± 0.11** | **-0.22 ± 0.04** | **-0.45 ± 0.13** | 0.09 ± 0.06 |

**Supplemental Figure 16: Phenotypic results of Barley lines.** Associated Barley lines of Barke, Bonus, Morex and HvLETM1-like2 mutated line FT880 with control and Taf treated plants. The log2 transformed and normalized to control phenotypes of Total Weight, Shoot Weight, Root Weight, Primary Root Length, Root Weight per Length, and Shoot to Root Ratio. Bars represent standard error. Asterisk represents significant difference between control and Taf treated total weight using an FDR *q* value of 0.05.

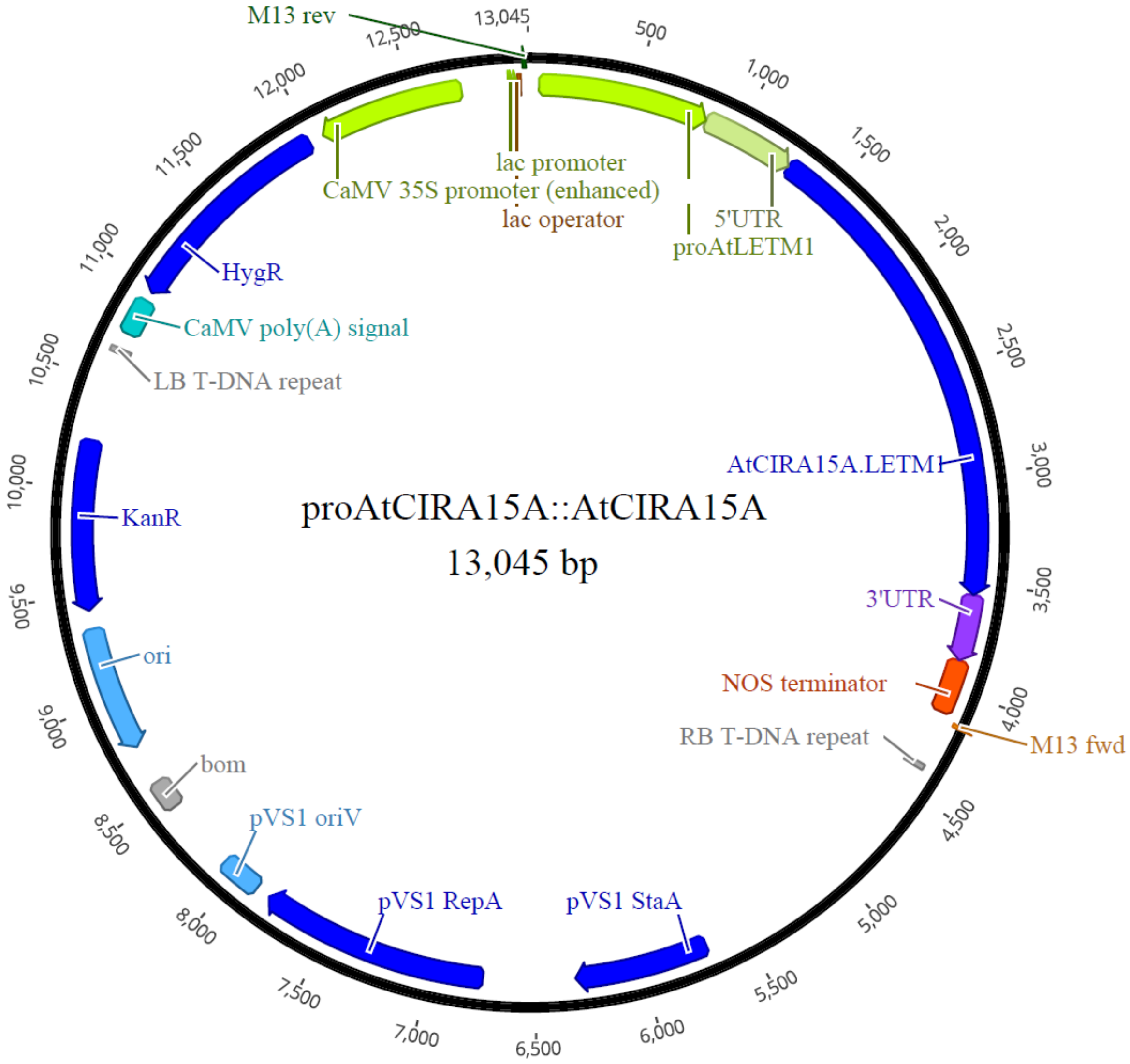


**Supplemental Figure 17: pro*AtCIRA15::AtCIRA15A/LETM1A* in pC1300 plasmid map**. Construct containing the synthesized product of *AtCIRA15A/LETM1A* promoter, 5'UTR and 3'UTR with the *AtLETM1A* CDS inserted. The backbone of the plasmid pC1300 gives hygromycin resistance under the constitutive 35S CaMV promoter and kanamycin resistance for agrobacterium transformation selection.

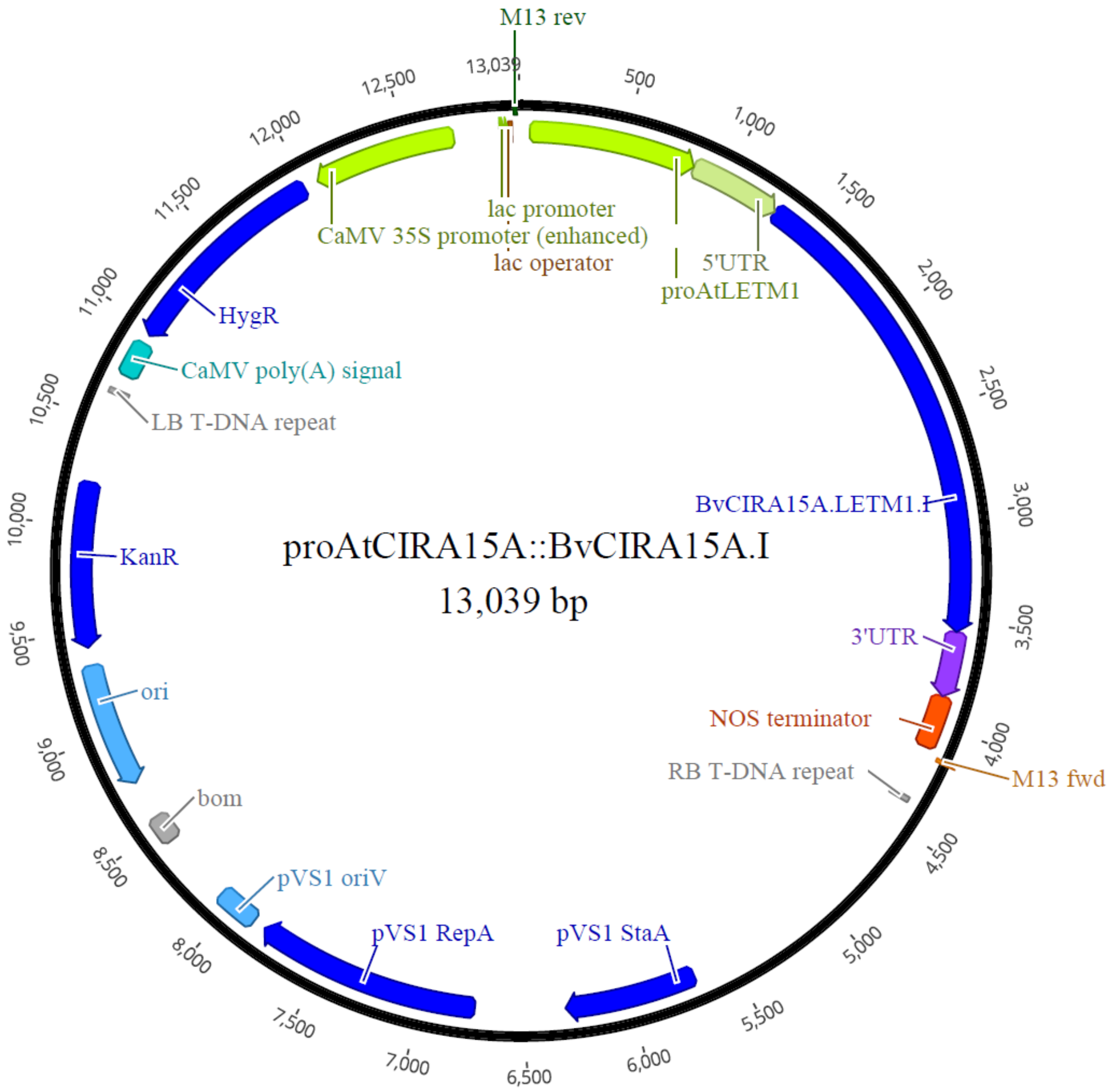


**Supplemental Figure 18: *AtCIRA15proBvCIRA15A/LETM1A.I* in pC1300 plasmid map.** Construct containing the synthesized product of *AtCIRA15A/LETM1A* promoter, 5'UTR and 3'UTR with the *BvLETM1A-I* allele CDS inserted. The backbone of the plasmid pC1300 gives hygromycin resistance under the constitutive 35S CaMV promoter and kanamycin resistance for agrobacterium transformation selection.

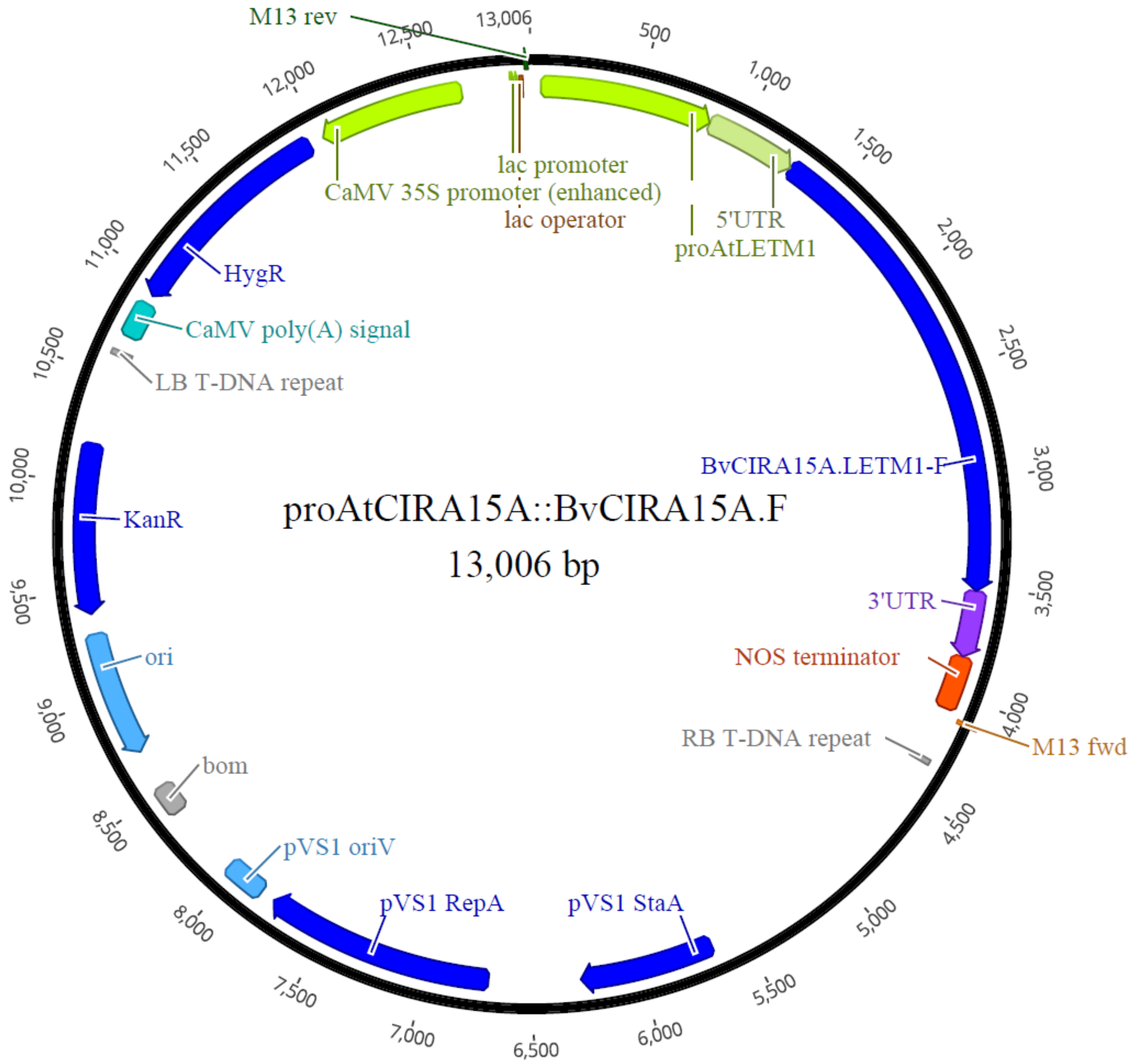


**Supplemental Figure 19: *AtCIRA15proBvCIRA15A/LETM1.F* in pC1300 plasmid map.** Construct containing the synthesized product of *AtCIRA15A/LETM1A* promoter, 5'UTR and 3'UTR with the *BvLETM1-F* allele CDS inserted. The backbone of the plasmid pC1300 gives hygromycin resistance under the constitutive 35S CaMV promoter and kanamycin resistance for agrobacterium transformation selection.

| Target | Forward Primer | Reverse Primer |
|---|---|---|
| **Construct verification** | | |
| *proAtCIRA15A::AtCIRA15A* | GGCCCTTGGGGTTCAAGTATA | GCAGGTCCTATCTTAAGAAC |
| | GGTTGGGGAGATTGGG | CAGCGGAGGGGTTGGA |
| *proAtCIRA15A::BvCIRA15A.I* | GGCCCTTGGGGTTCAAGTATA | AATAGGGAAGAAGGGATAG |
| | GGGCAGCCGAGGTGATG | CAGCGGAGGGGTTGGA |
| *proAtCIRA15A::BvCIRA15A.F* | GGCCCTTGGGGTTCAAGTATA | ATAAAAACTCCATGAAGGGG |
| | GGCAACCGAGGTGATG | CAGCGGAGGGGTTGGA |
| **AtCIRA15.LETM verification** | | |
| *cira15a-1* (Salk_042602C) | TCACAGGTTTAGTTTCCACGG | GGAGGGAATGGATACAAAAAG |
| *cira15a-2* (Salk_067558C) | AAGGATGAACACTGCAAATGG | CGTCGCTTCATTTTTCAACTC |
| *cira15b-1* (Salk_068877) | CATCTTTTTGGCTTGAAGCAG | TTTTCTTGTCTCCAATGGCTG |
| *cira15b-2* (Salk_051134C) | TACATCGCAGCCAATGCAAC | TTTCCGCGTATCATGGATCTTG |
| **BvCIRA15A.LETM1 allele verification** | | |
| *BvCIRA15A.LETM1* | AATGTAACGCCCAGGTTCTG | ACATAGGACTGTGTAGCAGGC |
| **HvLEMT1-like2 allele verification** | | |
| *HvLETM1-like2* | TGTCGTCTCGTCTGAACCAC | GCCTCCTCTCTAGCAGCCATG |

**Supplemental Figure 20: List of primers used in this paper.**
